# Structural deformations of lipid bilayers around membrane proteins inform oligomerization type

**DOI:** 10.64898/2026.11.24.769041

**Authors:** Gladys Díaz Vázquez, Samson G. F. Condon, Janice L. Robertson

## Abstract

A functional definition of a membrane protein’s structure is the three-dimensional arrangement of the polypeptide and the structure of the membrane around it. Biological membranes are composed of self-assembled lipid bilayers, with membrane proteins acting as inclusions that perturb this phase. These inclusions induce long-range changes in membrane structure and create an intrinsic, lipid-mediated driving force for membrane protein association. The question is how much different membrane protein folds shape their surrounding membrane to tune the affinity and specificity of this driving force. To investigate this, we conducted a computational study to evaluate whether different membrane protein oligomers exhibit characteristic membrane defects that indicate specificity in complexation. We performed coarse-grained molecular dynamics simulations of monomeric and oligomeric states of ten different channels and transporters in POPC bilayers and evaluated changes in membrane thickness, deformation energy, and lipid packing parameters. We found that all the proteins affect their surrounding membrane up to 3 nm from the protein surface. However, we observed larger defects at oligomeric interfaces for obligatory or structurally monomorphic complexes, suggesting that their oli-gomerization is partly driven by burial of membrane defects. In contrast, oligomers that adopt different quaternary structures across homologs show weaker perturbations at known binding interfaces, and larger perturbations at lipid-facing surfaces. Nevertheless, all systems show fewer membrane defects upon oligomerization, suggesting an underlying lipid-mediated driving force that each protein fold modulates to tune affinity via membrane-mediated oligomerization.

**Table of Contents (TOC) Graphic:** 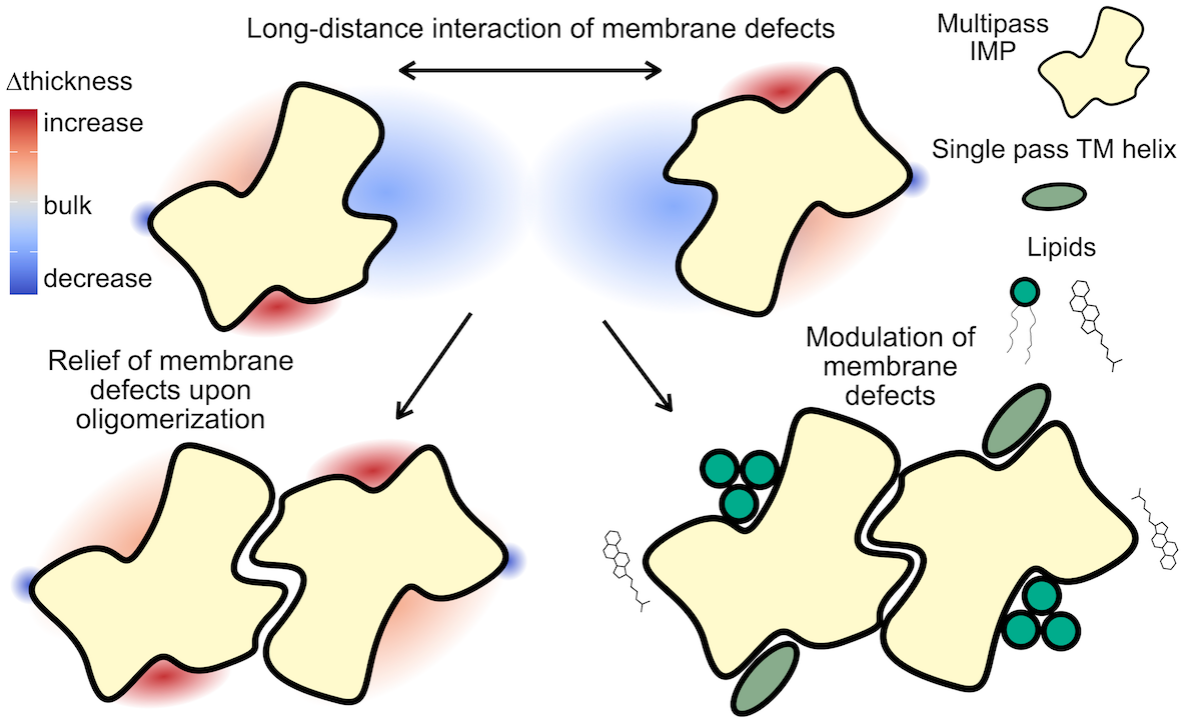

## INTRODUCTION

A protein’s structure is typically described solely by the arrangement of its polypeptide chain in three-dimensional space. This concept applies to soluble proteins, which are solvated in an isotropic, hydrophilic environment that lacks a continuous structure. The situation changes for integral membrane proteins (IMPs), whose main solvent self-assembles into a lipid bilayer with both hydrophilic and hydrophobic regions^1^. Notably, the membrane structure is not fixed but emerges as a thermodynamically favored phase from the highly dynamic lipid molecules. As such, membrane proteins have long been recognized as affecting the behavior of surrounding lipids and, in turn, influencing the membrane’s structure. Electron spin resonance (ESR) and nuclear magnetic resonance (NMR) studies showed that lipids in the annular shell, i.e., those directly associating with the protein, exhibit different dynamic behavior than lipids in the bulk phase^2^. These changes affect lipid packing, extending up to 2-3 layers beyond the annulus^3^, translating into long-range changes in the membrane’s 3-dimensional shape around an IMP^4^. Mean-field models predict that proteins impact the surrounding membrane up to 2.5 nm away, and that these perturbations introduce an intrinsic lipid-mediated driving force for protein aggregation^3,5–7^. Therefore, for IMPs, the protein conveys only part of the picture. Instead, a full functional depiction of a membrane protein’s structure requires the protein plus the surrounding membrane.

Experimental studies identifying long-range changes in membrane structure around membrane proteins have been challenging because lipids are highly dynamic. However, X-ray crystallography and cryo-EM studies have imaged lipids in the first annular shell that are in direct contact with the protein. Because these experiments are typically conducted under freezing conditions, and the protein constrains lipid configurations, these lipids become increasingly ordered and structurally resolvable. Usually, this picks out only a few of the most ordered lipids, but one study using X-ray solvent contrast modulation resolved the entire annular shell around the Ca^2+^ ATPase^8^. This revealed that phospholipid headgroup positioning is defined by tryptophans, arginines, and lysines, which perturb the typical bilayer structure. This results in an annular shell with local distortions that modulate bilayer thickness around the protein. These local changes are predicted to induce longer-range structural changes as the lipids optimize packing and other thermodynamic factors to achieve their most energetically favorable structural ensemble.

Recent cryo-electron microscopy (EM) and cryo-electron tomography (ET) studies of membrane proteins are now revealing long-range structural changes as well. Researchers first observed macroscopic distortions in detergent micelles and nanodiscs^9^, but recent innovations in liposome imaging show that these distortions also occur in lipid bilayers. For example, the large mechanosensitive ion channel Piezo perturbs the surrounding membrane, forming a dome-like structure in which force-induced membrane deformations activate the channel^10,11^. The TMEM16F scramblase locally thins the membrane near the protein, promoting lipid flipping between leaflets^12^. Notably, cryo-ET of reconstituted *F*_*1*_*F*_*O*_ ATP synthase shows that the protein dimerizes and then tends to self-assemble into rows, directing membrane ultrastructure into tubes and flattened discs, as observed in native mitochondrial cristae^13–15^. This does not rely on specific protein-protein contacts, but occurs to minimize membrane energetics^13,16^. Structural studies confirm that membrane proteins impart long-range changes in membrane shape, with dramatic impacts on biological function.

To better understand how membrane perturbations affect higher-order organization, we need more information on how different membrane proteins shape their surrounding membranes, and impact lipid configurations around them. Here, structural experiments fall short because lipids beyond the annular shell become unresolvable, and dynamic lipid configurations average out these signals. To fill this gap, theoretical modeling of membrane structure has been crucial. Continuum modeling and molecular dynamics simulations, both coarse-grained and all-atom, can predict large-scale structural changes in membranes^17–19^. Importantly, despite differing levels of chemical detail, provided sufficient sampling is obtained, results from these different methods converge, indicating that a protein’s effect on the membrane is robust. While each method offers insight into energetic changes, molecular simulations provide the information needed on lipid orientations and configurations related to protein-specific membrane changes.

This level of molecular detail has proven essential for understanding the driving forces behind large membrane proteins’ self-assembly into oligomers in membranes. Computational studies of the homodimeric CLC-ec1 chloride/proton antiporter and the fluoride channel Fluc-Bpe reveal marked membrane defects at the dimerization interface when the monomer is exposed^20,21^. Each hydrophobically mismatched surface causes local membrane thinning, inducing significant lipid tilting and disrupting lipid packing. Dimerization abolishes these defects, yielding a favorable dimerization free energy that experimental studies have shown to carry a thermodynamic signature of the hydrophobic effect^22^. In this case, however, it is more accurately described as the solvophobic effect, as it arises mainly from changes in lipid energetics. Notably, this mechanism, referred to as *burial of membrane defects*, drives orientational specificity for these oligomers: subunits first sense each other’s membrane defect, which funnels them along states that minimize membrane perturbations^20,22,23^.

If burying membrane defects is a generalizable driving force for membrane protein assembly, then we hypothesize that the size and shape of membrane defects co-evolve with specific types of membrane protein oligomers. To evaluate this, we studied ten different oligomeric ion channels and transporters with different assembly characteristics and performed coarse-grained molecular dynamics simulations to analyze changes in membrane shape and lipid configurations in monomeric vs. oligomeric forms. All the proteins studied perturb the membrane up to 3 nm from the protein surface. However, we observe larger defects at the association interfaces of defined and obligatory oligomers, whereas oligomers that associate in a variety of orientations across homologs show smaller deformations at the known binding interfaces than at lipid-facing regions. This suggests that membrane structure encodes information about oligomerization type and evolves with the membrane protein fold.

## RESULTS

### A structural comparison of channel and transporter oli-gomerization interfaces

To understand the nature of oligomeric interfaces and whether they uniquely impact their membranes, we examined a selection of IMPs from channels and transporters that self-assemble into oligomeric forms^24–33^. We organized these structures into three groups based on their assembly mode (**Fig. 1A**). First are the *obligate* oligomers, in which the association axis forms the functional pathway (e.g., an ion-conducting pore or substrate pathway). In this case, all subunits are required for function, typically conferring residue interactions that simultaneously co-ordinate ions or substrates. In this group, we include the 4-trans-membrane (TM) helix inverted topology homodimeric fluoride channel Fluc-Bpe^31^, the 3-TM helix homodimeric sugar transporter SemiSWEET^29^, and the 2-TM helix homotetrameric potassium channel KcsA^33^.

**Figure 1.**
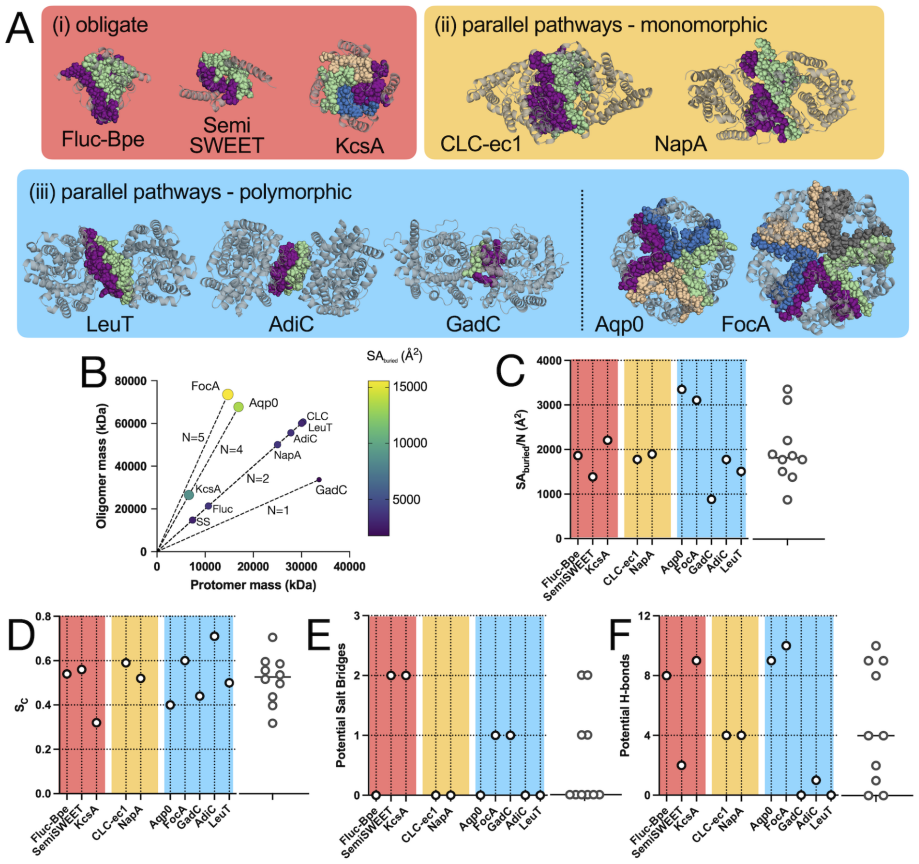
Physicochemical analysis of channel and transporter oligomerization interfaces. Ten channels and transporters from three oligomerization groups: (i) obligate oligomers (red): Fluc-Bpe (PDB 5NKQ), Semi-SWEET (PDB 4X5N), and KcsA (PDB 1K4C); and two types of parallel-pathway oligomers: (ii) monomorphic (yellow): CLC-ec1 (PDB 1OTS), NapA (PDB 5BZ2), and (iii) polymorphic (blue): LeuT (PDB 6XWM), AdiC (PDB 7O82), GadC (PDB 4DJK), Aqp0 (PDB 2B6P), FocA. (PDB 3KCU). (B) Total buried surface area for each oligomer, *SA*_*buried*_ (Å^2^). (C) Buried surface area normalized by subunit stoichiometry, *N*. (D) Shape complementarity co-efficient, *S*_*C*_. (E) Potential salt-bridge and (F) hydrogenbond pairs per subunit at each oligomeric interface. C-F: plots on the right show combined distributions. Calculations were carried out with PDBePISA.

Next are the parallel-pathway oligomers, in which each subunit contains a distinct permeation or transport pathway. Two types are observed. The first is defined as *monomorphic* because only a single oligomeric interface has been observed for that protein family’s fold, suggesting highly specific assembly at that interface. Examples include the 18-TM helix homodimeric CLC-ec1 Cl^−^/H^+^ antiporter^25^ and the 13-TM helix homodimeric Na^+^/H^+^ antiporter NapA^24^. Unlike obligate complexes, the role of oligomerization is unclear; typically, oligomerization is not required for function but could play a regulatory role.

We call the second type of parallel-pathways oligomers *polymorphic* because subunits of the same fold can assemble at different interfaces, resulting in different quaternary arrangements. Researchers have observed this assembly type in multiple protein families, particularly the Amino Acid-Polyamine Organocation (APC) superfamily, which adopts a “LeuT-fold” comprising 10 TM helices, with 1 or 2 additional TM helices on the N- or C-terminal side^34^. Despite the common fold, these transporters adopt various oligomeric arrangements, monomers, dimers, and trimers, requiring association at different interfaces. The glutamate-GABA antiporter GadC is believed to be a monomer; however, it crystallized as an inverted-topology dimer via helices 11 and 12, likely an artifact^30^. On the other hand, the homologous L-arginine/agmatine antiporter AdiC forms a defined homodimer via TMs 11 & 12^28^. In contrast, the Na^+^/leucine transporter LeuT has been structurally resolved as monomers^35^ and dimers, but here it assembles at a different interface than AdiC, via TMs 9 & 12^27^. Another superfamily that exhibits polymorphic oligomerization is the Major Intrinsic Protein (MIP) superfamily, which contains 6 TM helices that form an hourglass fold. Of these, the water channel Aqp0^26^ forms tetramers, whereas proteins from the Formate-Nitrate Transporter (FNT) subfamily, e.g., FocA, adopt pentameric arrangements^32^, despite homologous subunit structure. Subunits adopting multiple oligomeric forms despite similar structures raise questions about the factors that determine oligomerization preferences across homologs.

For these ten oligomers, we analyzed the physico-chemical properties of the interaction interfaces of the membrane-embedded components (**Fig. 1, Table 1 & Supp. Table 1**). The transmembrane protomer masses range from 6.6 to 33.7 kDa (**Fig. 1B**). The buried surface area per subunit ranges from 880 to 3350 Å^2^, with dimers having a narrower distribution than higher-order oligomers. For dimers, the buried surface area per subunit falls within a narrow distribution of 1584 ± 363 Å^2^ (mean ± std), whereas for higher-order oligomers it jumps to 2888 ± 602 Å^2^ (**Fig. 1C**). This is because higher-order packing involves two interfaces that form a V-like shape (**Fig. 1A**).

**Table 1.**
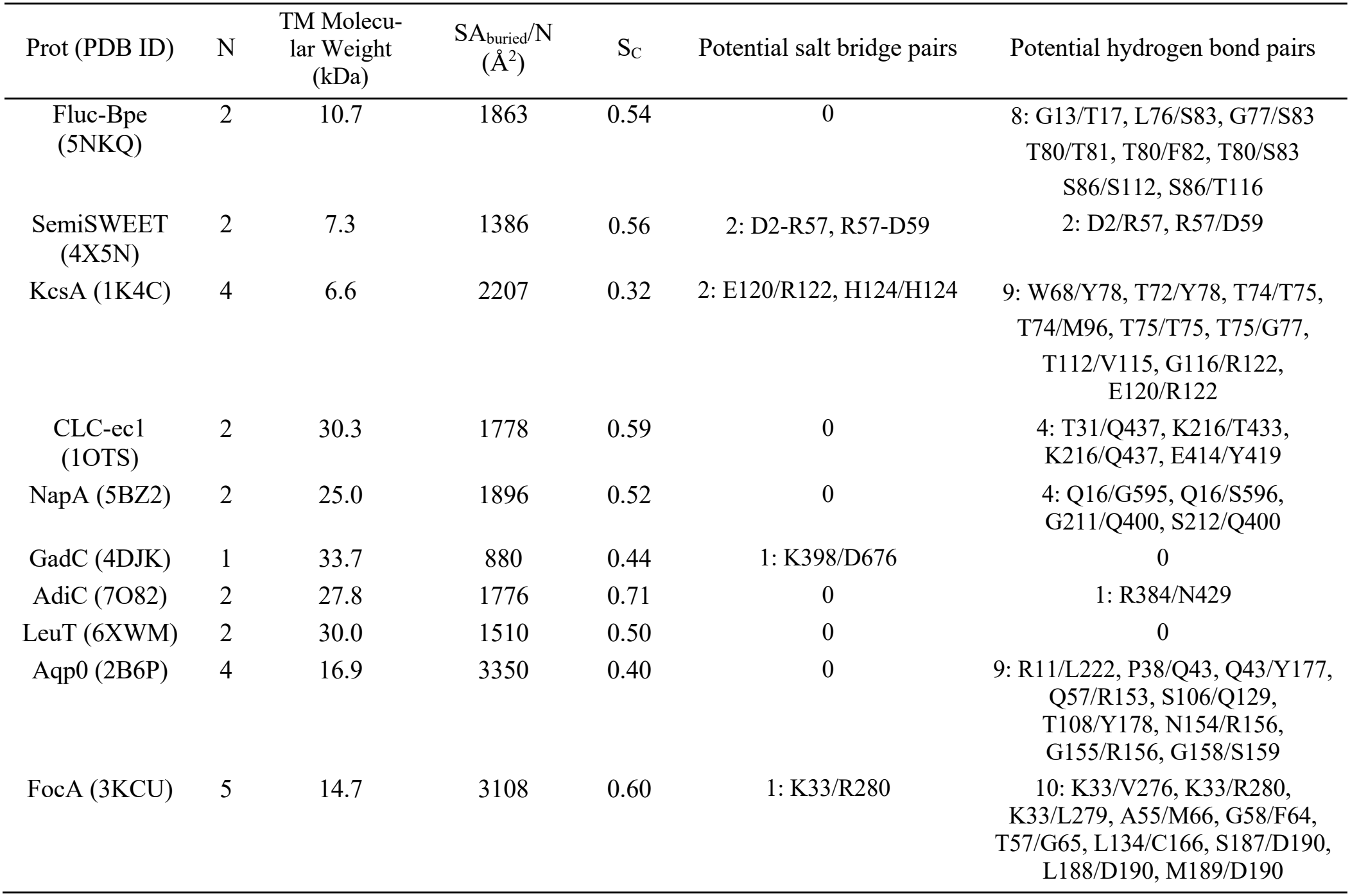
Surface complementarity, salt bridges & hydrogen bonds of membrane protein oligomers.

| Prot (PDB ID) | N | TM Molecu-lar Weight (kDa) | SA<sub>buried</sub>/N (Å<sup>2</sup>) | S<sub>C</sub> | Potential salt bridge pairs | Potential hydrogen bond pairs |
|---|---|---|---|---|---|---|
| Fluc-Bpe (5NKQ) | 2 | 10.7 | 1863 | 0.54 | 0 | 8: G13/T17, L76/S83, G77/S83<br>T80/T81, T80/F82, T80/S83<br>S86/S112, S86/T116 |
| SemiSWEET (4X5N) | 2 | 7.3 | 1386 | 0.56 | 2: D2-R57, R57-D59 | 2: D2/R57, R57/D59 |
| KcsA (1K4C) | 4 | 6.6 | 2207 | 0.32 | 2: E120/R122, H124/H124 | 9: W68/Y78, T72/Y78, T74/T75, T74/M96, T75/T75, T75/G77, T112/V115, G116/R122, E120/R122 |
| CLC-ec1 (1OTS) | 2 | 30.3 | 1778 | 0.59 | 0 | 4: T31/Q437, K216/T433, K216/Q437, E414/Y419 |
| NapA (5BZ2) | 2 | 25.0 | 1896 | 0.52 | 0 | 4: Q16/G595, Q16/S596, G211/Q400, S212/Q400 |
| GadC (4DJK) | 1 | 33.7 | 880 | 0.44 | 1: K398/D676 | 0 |
| AdiC (7O82) | 2 | 27.8 | 1776 | 0.71 | 0 | 1: R384/N429 |
| LeuT (6XWM) | 2 | 30.0 | 1510 | 0.50 | 0 | 0 |
| Aqp0 (2B6P) | 4 | 16.9 | 3350 | 0.40 | 0 | 9: R11/L222, P38/Q43, Q43/Y177, Q57/R153, S106/Q129, T108/Y178, N154/R156, G155/R156, G158/S159 |
| FocA (3KCU) | 5 | 14.7 | 3108 | 0.60 | 1: K33/R280 | 10: K33/V276, K33/R280, K33/L279, A55/M66, G58/F64, T57/G65, L134/C166, S187/D190, L188/D190, M189/D190 |

The shape complementarity coefficient, *S*_*C*_, quantifies the degree to which two surfaces fit together and ranges from 0-1, with 1 indicating a perfect fit^36^. For reference, anti-body-antigen complexes are generally in the range of 0.64-0.68, while soluble protein-protein complexes can be higher, around 0.70-0.76^36^. The *S*_*C*_ coefficients for these membrane-embedded complexes are lower. Excluding the extremes of KcsA (0.32) and AdiC (0.71), the rest of the examples fall within a range of 0.4-0.6, with a 0.52 ± 0.07 (mean ± std) (**Fig. 1D**). This indicates that the membrane-embedded oligomers exhibit complementarity, but not as strongly as observed for soluble protein binding complexes. Notably, we see no differences among the oligomer types, and the GadC dimer, thought to be artifactual due to crystallization conditions, shows greater surface complementarity than KcsA or Aqp0, even though it does not represent an evolved binding interface.

We also identified potential charged and polar protein interactions at these membrane-embedded interfaces. Salt bridges can strongly stabilize an oligomeric assembly because exposing free charges to the lipid bilayer is highly unfavorable. However, salt bridges are rare, with only obligate oligomers showing more than one potential salt bridge (**Fig. 1E**). Hydrogen-bonding pairs are more likely to occur, especially in the larger complexes like Aqp0 and FocA, but are rare in other systems, such as the LeuT-fold transporters (**Fig. 1F**).

Across these protein interfaces, we do not see outstanding trends in oligomerization behavior, particularly between the parallel assembly pathways sub-types. Even the non-native GadC dimer shows comparable metrics to evolved oligomers, indicating that protein structure alone makes it challenging to infer membrane protein oligomerization. This suggests that a deeper investigation into the surrounding membrane is warranted for a more complete understanding of the determinants for oligomerization.

### The impact of a membrane protein on its surrounding membrane

We first investigated how a fully assembled oligomeric complex affects its surrounding membrane using molecular dynamics (MD) simulations. Following well-validated protocols^20,21,23^, we used the coarse-grained MARTINI 2.2 force field^37^ to simulate the large-scale membrane structural changes around a specific conformation of each membrane protein complex (**Supp. Fig. 1 & 2**). Note that simulations with this coarse-grained force field have been shown to converge with long-timescale all-atom MD simulations^21,23^, yielding similar results but on a practical computational timescale. In addition, we study each oligomer in a POPC bilayer, which we selected to facilitate proper sampling of the membrane and its structural changes. Multi-component lipid mixtures are notoriously difficult to converge, which can impede analysis of equilibrium lipid and membrane behavior^20,21,23^. Simulations were run for 10 μs, a timescale where lipid exchange sampling is observed to converge to > 95% in each system (**Supp. Fig. 3**).

Using the grid-based membrane analysis software suite MOSAICS^38^, we calculated a two-dimensional grid-based map for each system of the mean change in membrane thickness with respect to the bulk, 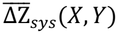 (**Fig. 2A**). Each oligomer shows changes in membrane thickness surrounding the protein, often as thinning right near the protein surface, followed by a wave of compensatory thickening, until it relaxes to the bulk membrane thickness. The deviations appear largely symmetric, except for GadC, where the dimer structure is expected to be in a non-native configuration. To quantify significant differences in membrane deformation, we calculated the Z-score for each grid point, describing how many standard deviations it is from the membrane thickness distribution of a POPC lipid bilayer without protein. To determine the Z-score threshold for significant changes in membrane, we applied a false discovery rate (FDR) approach set to 5% across all grid elements^39–42^. This analysis identified a Z-score threshold of 0.058, corresponding to a 0.014 nm change in mean membrane thickness as being significant (**Supp. Fig. 4**).

**Figure 2.**
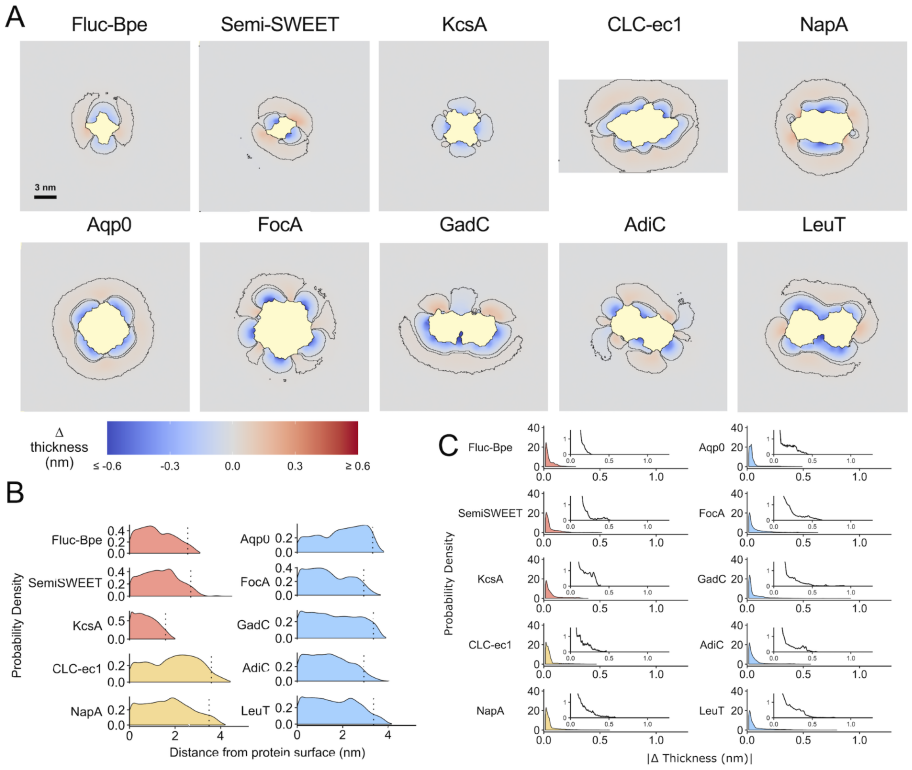
The impact of an oligomeric membrane protein on its surrounding membrane. (A) 2D membrane thickness maps for each oligomer from coarse-grained MD trajectories averaged over three independent runs. Black lines outline significantly deformed regions of the membrane identified from the FDR-based thresholding. (B)Probability density distributions of significantly perturbed grid points as a function of the distance from the surface of the protein. (C) Probability density distributions of significantly perturbed grid points as a function of the absolute magnitude of membrane thickness changes. For each plot, an inset is provided to magnify the long tail of the same distributions.

Analysis of the significantly perturbed regions shows how far deformations extend from the protein surface (**Supp. Fig. 5**). Probability density distributions of the perturbed grid elements plotted as a function of distance from the protein surface show that most systems affect their surrounding membrane up to 3-4 nm away from the protein surface. The exception is KcsA, where defects are more localized to within 2 nm (**Fig. 2B, Supp. Fig. 6**). Most of the deformations are small, within 1 Å, but each protein system exhibits regions where more extreme deformations are found, in the range of 2-5 Å or more (**Fig. 2C**). Overall, this shows that oligomeric complexes affect their surrounding membrane in small but significant ways up to 3 nm away.

### Monomorphic oligomers exhibit significant defects at oli-gomerization interfaces

Next, we investigated whether isolated subunits from these complexes exhibit characteristic defects at their oligomerization interfaces consistent with a driving force for association via burial of membrane defects. To test this, we calculated the mean change in membrane-thickness maps from CGMD simulations of isolated monomers (**Fig. 3A**). When isolated, monomers are no longer constrained by other subunits and adjust their alignment relative to the membrane normal. The subunits are generally oriented close to the membrane normal, starting at 4.8 ± 0.3° for CLC-ec1, but appear quite tilted in some systems: 17.3 ± 0.3° for Fluc-Bpe and 17.7 ± 1.2° for KcsA (**Fig. 3A, Supp. Table 2**). These two ion channel complexes form obligatory oligomers, with strongly charged binding interfaces because they form the aqueous pore that binds ions. As such, the tilt angles observed here could reflect a reorientation of the subunit such that polar/charged regions minimize exposure to the hydrophobic membrane core. Indeed, Fluc-Bpe and KcsA exhibit by far the largest change in tilt angle between the monomeric and oligomeric states, with Fluc-Bpe being 9.98 ± 2.25º and KcsA 11.42 ± 1.63º (**Supp. Table 3 & 4**).

**Figure 3:**
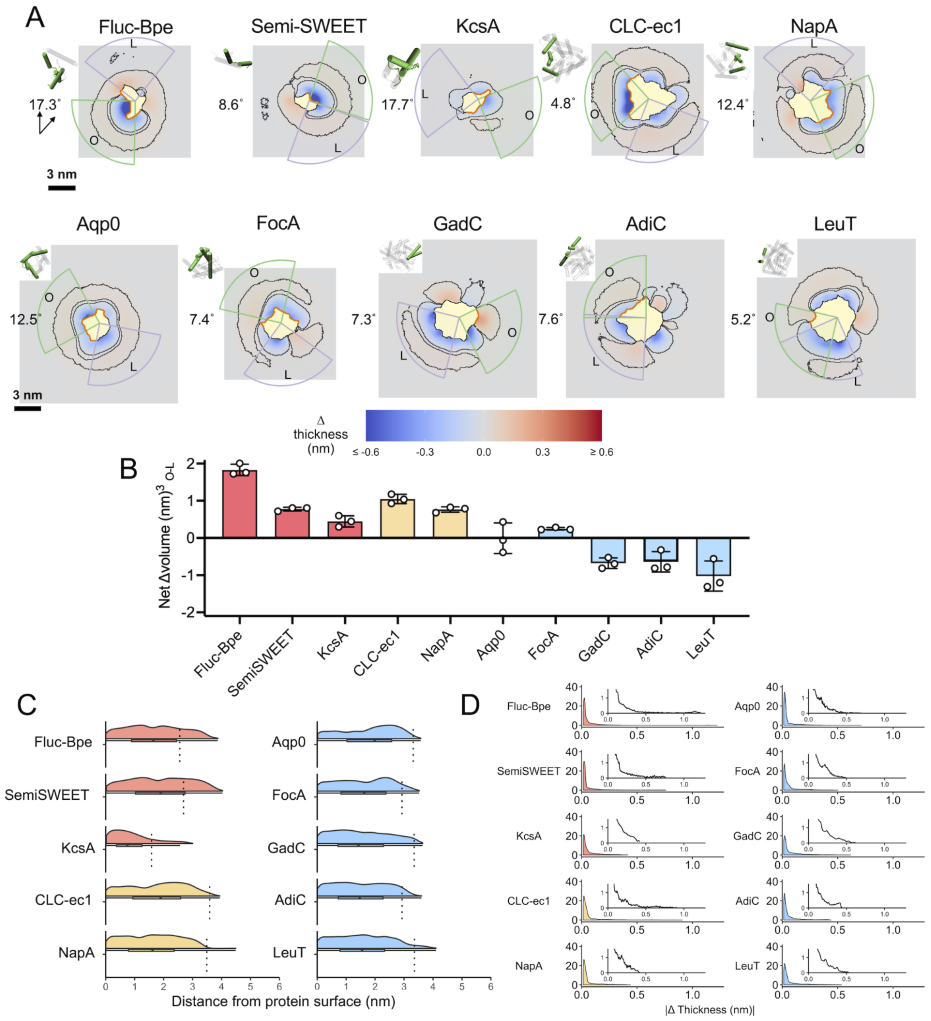
Membrane perturbations at buried vs. lipid-facing interfaces in monomeric subunits. (A) Average membrane thickness maps for each channel and transporter monomer (upper leaflet as target for calculation). Black lines outline significantly deformed regions of the membrane identified from the FDR-based thresholding. In each map, the green arc corresponds to the face of maximum perturbation that overlaps with the oligomeric interface (orange line), and the purple arc corresponds to the face of maximum perturbation that faces the lipids. (B) Bar plot showing the difference between the net deformed volume at the oligomeric (green) and lipid-facing faces (purple). (C)Probability density distributions of significantly perturbed grid points as a function of the distance from the surface of the protein. (D) Probability density distributions of significantly perturbed grid points as a function of the absolute magnitude of membrane thickness changes. For each plot, an inset is provided to magnify the long tail of the same distributions.

To assess whether the oligomeric interface of each protein displays the largest membrane defect, we carried out a rotating arc analysis around the protein, quantifying the sum of absolute membrane volume changes within the arcs (**Fig. 3A, Supp. Fig. 7 & 8**). This allowed for localization of the arc with the maximally perturbed membrane either at the oli-gomerization (O) or lipid-facing (L) protein face. Taking the difference allows us to examine whether an oligomeric interface is more perturbative (O-L > 0) or whether a larger defect occurs at a lipid-facing surface (O-L < 0). Among the systems studied, obligatory and monomorphic complexes show the largest deformation at the evolved oligomerization interface (**Fig. 3B, Supp. Fig. 7**). In contrast, polymorphic assemblies show either no difference or a larger membrane defect at a lipid-facing region. Aqp0 and FocA show deformations at the oligomerization interface similar in magnitude to those facing the membrane. However, LeuT-fold transporters GadC, AdiC, and LeuT all show larger perturbations at lipid-facing faces. This highlights a correlation between large membrane defects at the oligomerization interface and oligomers that evolved defined binding structures.

The distribution of distances of significantly perturbed grid elements (**Fig. 3C, Supp. Fig. 6**) is like that in the assembled oligomers (**Fig. 2C**), indicating that the monomers affect a similar area of the surrounding membrane. However, when examining the magnitude of the deformations (**Fig. 3D**), the obligatory and monomorphic complexes Fluc-Bpe, Semi-SWEET, and CLC-ec1 have localized defects with much more extreme changes in thickness than the other proteins.

### Membrane perturbations are minimized upon oligomerization

Burial of membrane defects and reduced hydrophobic mismatch have been energetically linked to the equilibrium stability of membrane protein oligomerization^13,16,22,23,43^ (**Fig. 4A**). This suggests that the state-dependent changes in membrane perturbations observed could drive oligomeric assembly. To predict which complexes would be strongly driven by burial of membrane defects, we calculated the change in membrane deformation volume upon oligomerization for each of the systems in their native assembly forms. The absolute total deformation was calculated by summing over all significantly perturbed grid elements in the monomer and multiplying by *n* (the number of subunits in the oligomeric complex) **(Fig. 4B)**, and for the intact oligomer system) 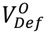 **(Fig. 4C)**. The net change in volume defects upon oligomerization is calculated as: 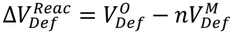 **(Fig. 4D)**. For all complexes studied, we find a net decrease in membrane perturbation upon oligomerization, 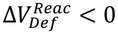, suggesting that defects generally decrease as subunits associate (**Fig. 4D**). This means that the membrane maximizes the area that is in a bulk lipid bilayer state upon oligomerization in all systems.

**Figure 4:**
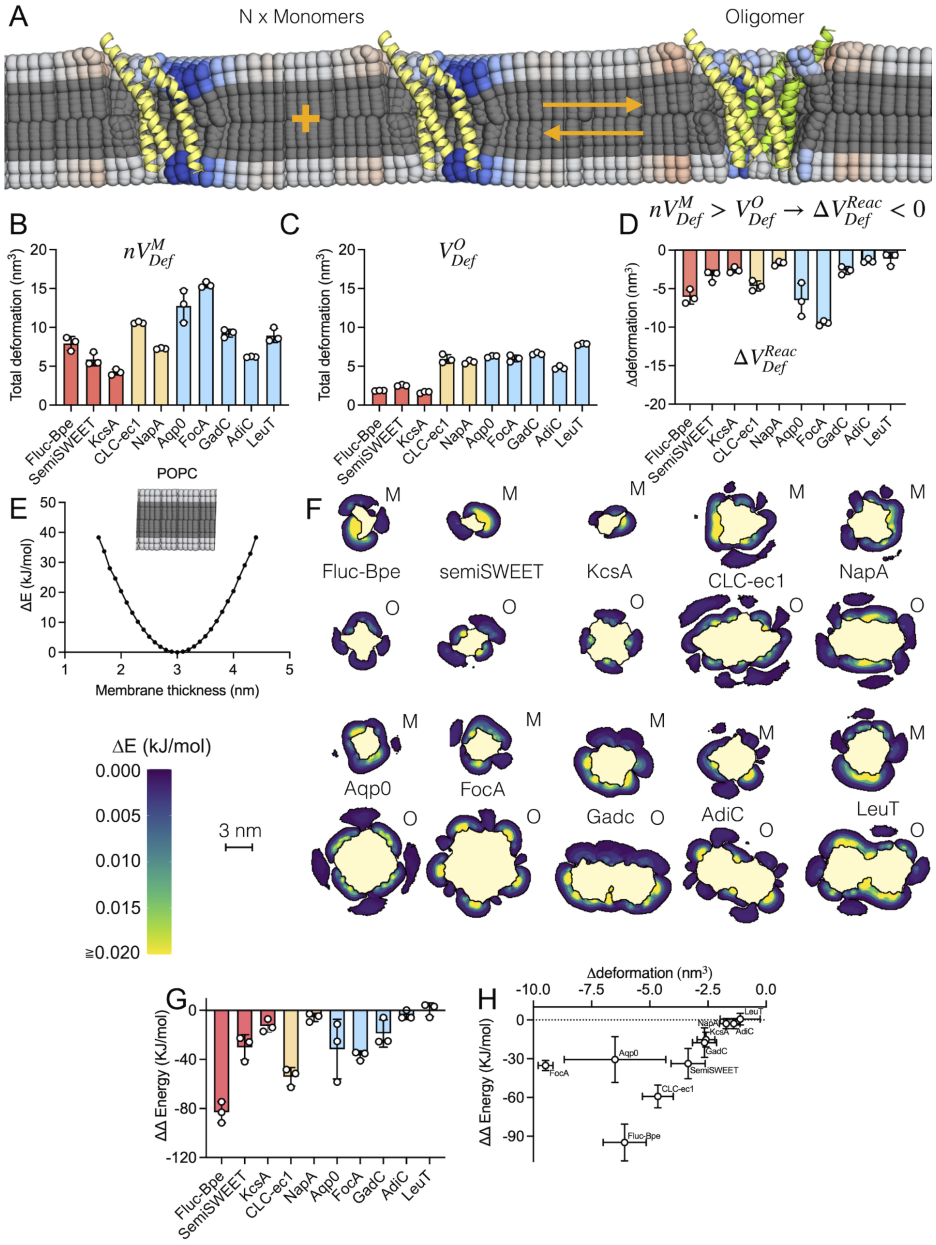
Homo-oligomerization is accompanied by relief of membrane defects. (A) Representation of the homo-oligomerization of an integral membrane protein and a change in lipid membrane thickness. In this representation, the thinning at the oligomeric interface (blue lipid head-groups) is eliminated by protein-protein interactions, but the deformations on the other side of each monomer (red lipid head-groups) remain. In total, 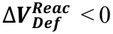 for this system. (B-D) From left to right, net volume of membrane deformations for the separated monomers 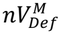 (B), net volume of membrane deformations for the oligomeric state 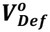 (C), and the change in the net deformation volume accompanying the oligomerization reaction 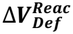(D). (E) Thickness-dependent potential-of-mean-force function derived from a simulation of a purely POPC bilayer. (F) Average membrane thickness energy maps (upper as target leaflet for calculation) for both monomer (M) and oligomer (O) states. (G) Change in energy between the oligomeric state and the separated monomers. (H) Change in Estimated energy vs change in net deformed volume between monomer and oligomer states.

Although defects become buried upon oligomerization in every system, the magnitude of the volume change varies by system, suggesting that the membrane confers differential stability. The largest volume changes are observed for the higher-order complexes FocA & Aqp0, which have the most to gain from minimizing membrane defects because of the substantial number of subunits in the assembly. This is followed by the obligatory and monomorphic complexes Fluc-Bpe ~ CLC-ec1. The remaining systems fall within a similar range of membrane-defect minimization, suggesting a non-specific, but minimal stabilizing effect arising from the membrane, at least for the states studied here.

However, the energy change from burying membrane defects depends not only on the total perturbed volume, but also on the magnitude of local perturbations. Stronger defects are much more energetically costly than many small defects. To assess this, we calculated the free-energy cost by mapping the membrane deformations onto the thickness-dependent potential-of-mean-force energy function of a pure POPC lipid bilayer (**Fig. 4E**). Note that this represents only one component of the actual free energy change and does not account for contributions from the protein, including membrane-protein interactions. Still, this allows us to examine the membrane-only contribution and generate 2D free-energy maps for each system in both the monomer and oligomer states (**Fig. 4F**). Taking the difference as before, we obtain the change in free energy upon oligomerization from burial of membrane defects (**Fig. 4G, Supp. Fig. 9**). The energy differences change the ranking as follows: Fluc-Bpe > CLC-ec1 > FocA > Aqp0 > SemiSWEET > all others. In all systems, many of the grid points with significant changes in membrane thickness have no associated energetic penalty, restricting the range of energetically unfavorable perturbations **(Supp. Fig. 10)**. However, apart from FocA and Aqp0, there is generally a strong, monotonic relationship between the change in net deformed membrane volume and the estimated difference in membrane energetics between the monomeric and oligomeric states **(Fig. 4H)**. The deviation from the trend for FocA and Aqp0 appears to reflect the number of grid points with a significant change in membrane thickness but no associated energetic penalty, combined with the higher stoichiometry for these two proteins (**Supp. Fig. 5, Supp. Fig. 10**). In this analysis, NapA, AdiC, and LeuT show no favorable energy change upon oligomerization. However, this does not mean they are unstable oligomers; rather, the membrane is not a major driving force for assembly at the oligomeric interface studied.

### Characteristic changes in lipid configurations at defects

Membrane defects may not always be reflected in membrane thickness changes. Protein interfaces can also introduce packing defects, introducing significant energetic penalties for solvation while maintaining the same lipid bilayer thickness. To assess whether packing defects occur characteristically at obligatory and monomorphic oligomerization interfaces vs. polymorphic interfaces, we analyzed the lipid configurational changes using MOSAICS (**Fig. 5, Supp. Fig. 11**). We performed separate analyses of the upper (U) and lower (L) leaflets of the bilayer for changes in four lipid metrics relative to the bulk: lipid acyl-chain length (**Fig. 5A**), lipid tilt angles (**Fig. 5B**), lipid interdigitation changes (**Fig. 5C**), and water-chain contacts (**Fig. 5D**). In general, we see three systems standing out in their impact on altering lipid configurations at the oligomerization interface compared to the bulk – CLC-ec1, NapA, and Fluc-Bpe, which are all monomorphic or obligatory oligomers. NapA did not show a significant membrane thickness defect, but it stands out as producing a packing defect that could also drive oligomerization at this interface.

**Figure 5:**
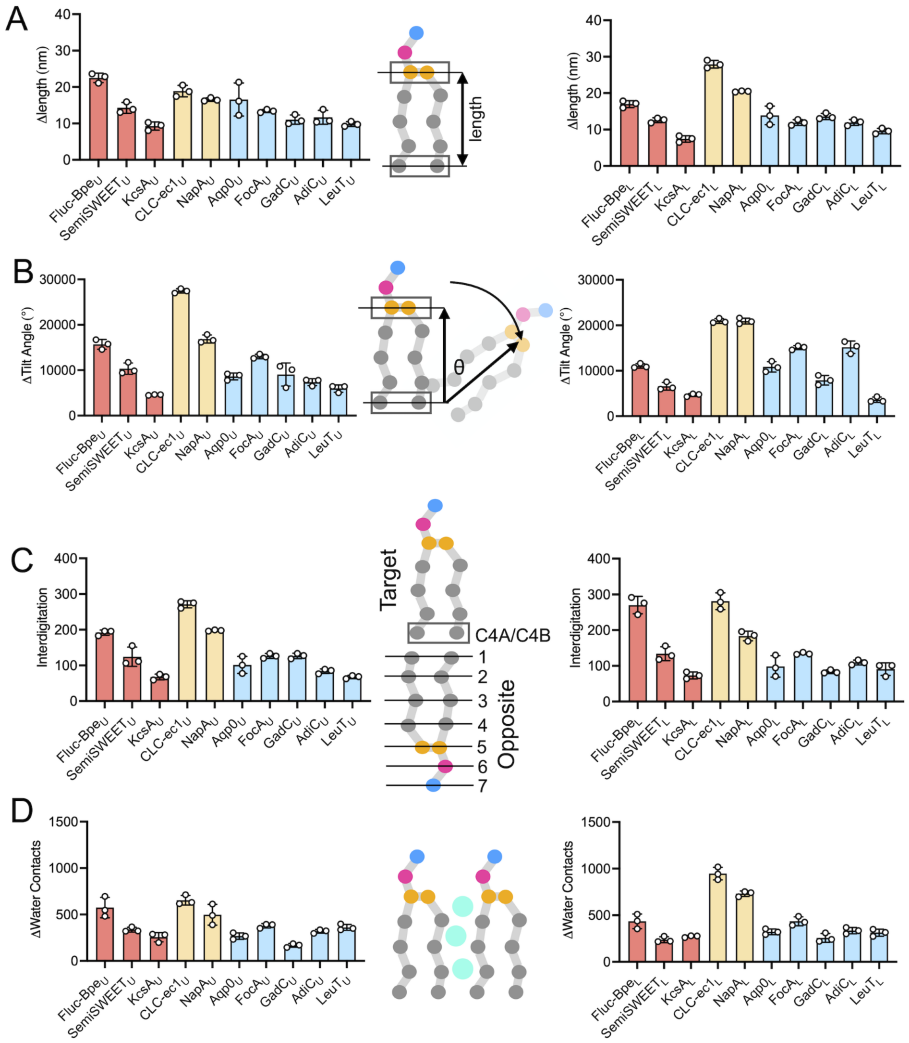
Summations of perturbations of lipid configurations in the upper (left) and lower (right) leaflets. (A) Lipid length, defined as the distance between the selected beads (boxed). (B) Lipid tilt angle, defined as the angle between the membrane normal and a vector defined by the selected beads (boxed) (C) Lipid interdigitation is measured by defining beads C4A/C4B in the target leaflet and measuring its contact with the ranked beads on the opposite leaflet. (D) Lipid-water contact is measured by counting the number of water molecules that enter in contact with the hydrophobic region of the lipids (gray beads). For each analysis from A-D the data points represent the sum of perturbations within the maximal 45° arc overlapping the oli-gomeric interface.

We used a rotating arc analysis to identify regions of maximal deviation in each of these lipid characteristics (**Supp. Fig. 11**). As with membrane thickness, obligatory and monomorphic oligomer subtypes showed larger deviations at the oligomerization interface than in lipid-facing regions. Interestingly, this trend diverges in the polymorphic oligomers, with Aqp0/FocA hovering around no net change and GadC/AdiC/LeuT showing larger defects at the lipid-facing interfaces. This identifies configurational membrane perturbations that may be energetically significant but are not buried upon oligomerization, raising the possibility of other functional roles of membrane defects, such as substrate binding, lipid interactions, or interactions with other protein partners.

## DISCUSSION

Since the earliest conceptualization of membrane proteins in lipid bilayers, researchers have recognized that while proteins can be stable within membranes, they likely perturb the surrounding membrane structure and energetics. Mean-field theoretical models suggested that proteins perturb the membrane up to 2-3 shells beyond the annular lipids^4,6,7^, imposing an implicit driving force for assembly. If true, this suggests that membrane protein folds and oligomeric assemblies evolved in the context of this physical driving force, producing characteristic changes in membrane perturbations.

Examining this in molecular detail for a subset of oligomeric complexes, we observe several findings that support this. First, obligatory and monomorphic oligomers show their maximum membrane perturbation at the oligomerization interface. Polymorphic complexes, on the other hand, do not. Second, systems that gain the most energetic benefit from burying the membrane defect fall into the obligatory or monomorphic categories, i.e., Fluc-Bpe & CLC-ec1, respectively. Finally, the largest lipid configurational defects occur in the monomorphic assemblies CLC-ec1 and NapA. In these three ways, we find characteristic membrane defects, largely in membrane thickness, but also lipid configurations associated with native binding interfaces that exhibit orientational specificity and strong affinity. This suggests that the membrane protein’s fold may have co-evolved to use the driving force arising from the membrane.

Another finding is that all proteins perturb their surrounding membranes to similar distances, up to 3-4 nm, corresponding to the 2-3 lipid shells predicted by mean-field theory. This applies to both the assembled oligomers and the isolated monomers, and defects arise from essentially all protein surfaces, whether protein-facing or lipid-facing. All systems show a net relief in deformed membrane volume for homo-oligomerization. This is small for the polymorphic systems, but it adds up for the higher-order oligomers Aqp0 and FocA, suggesting a substantial membrane-driven driving force for assembly, even when membrane defects are small. This likely reflects a non-specific driving force for membrane protein assembly into higher-order states such as 2D crystalline arrays. For example, in Aqp0, membrane-thickness defects at the lipid-facing interfaces appear comparable to the defect at the oligomerization interface, suggesting they could guide higher-order organization of Aqp0 tetramers in membrane.

This may also explain the structural biology observation that many membrane protein oligomers are expected to be in non-native states. For example, we studied the GadC dimer here, which is expected to be a crystallographic artifact because the subunits form an antiparallel dimer, with no evidence that it exists in the biological environment. Because burial of membrane defects is expected to occur in other hydrophobic solvent environments, such as detergent micelles, we expect GadC to associate during purification to hide a perturbative interface, and this orientation is likely favored over other possibilities due to additional protein interactions. This likely applies to many other systems that have been shown to form non-native assemblies^46^, supporting the idea that there is a nonspecific driving force for membrane protein assembly to minimize their surface area exposed to the surrounding hydrophobic solvent.

One interesting finding arose in our analysis of the polymorphic LeuT-fold dimers. Each system studied maintains a similar subunit fold, yet they dimerize at different interfaces (LeuT – TM9 & TM12, AdiC – TM11 & TM12, GadC). However, the maximum deformations always occurred at a lipid-facing region outside of the known oligomerization interface. The fact that this occurs even when the oli-gomerization interface itself is not well defined suggests that this protein fold evolved to incur multiple membrane defects. This aligns with these proteins’ oligomerization without a common orientational specificity, suggesting that protein interactions instead control orientation. Indeed, molecular dynamics simulations of BetP identified candidate salt-bridge interactions involved in trimer formation, with W101A/T351A mutations leading to a monomeric form of BetP^47^. Alternatively, membrane defects at multiple sites could suggest that there is not one defined oligomeric assembly, but that different orientations could occur as part of a regulatory mechanism. Finally, defects at these external faces may also create lipid-modulation sites or suggest interactions with other proteins for functional regulation. Interestingly, members of the monoamine transporter (MAT) family, such as SERT^48^ and DAT^49^, also adopt a LeuT-fold, regulate oli-gomerization using lipids like cholesterol and PIP_2_, and form multiple types of oligomeric rearrangements^50–52^. LeuT-fold proteins may have evolved to use lipids and other molecules to regulate oligomerization, facilitating their ability to acquire different oligomeric states in the cell.

In summary, membrane proteins alter the surrounding membrane structure, creating an implicit driving force for assembly that minimizes membrane perturbations. This has evolved alongside the protein structure, playing a role in defining oligomeric orientational specificity in obligatory and monomorphic complexes. We suspect that the extent of these defects contributes to oligomer stability, although overall affinity also depends on other factors, including protein-lipid and protein-protein interactions. Because the protein fold determines membrane shape, however, this membrane-dependent driving force is likely shared across entire protein families. This may also apply to the assembly of different membrane protein components, as in macromolecular assemblies, and to the non-specific assembly of protein substrates with membrane-embedded enzymes such as proteases or insertases. Importantly, because this driving force is entirely membrane-dependent, it is tunable by changes in lipid composition^20^, high-lighting how membrane chemistry may shape membrane protein ultrastructure. In each case, burial of membrane defects likely brings components close enough to sample and explore fine protein chemistry. A fuller understanding will require further computational studies of membrane defects around proteins, as well as experimental studies that quantify changes in stability and specificity.

## METHODS

### Preparation of protein-membrane systems

The PDB files used in this study are listed in **Table 1**. When necessary, MODELLER version 10.6^53^ was used to rebuild missing loops in the original PDBs, and the resulting structure file was used in the simulations. The CHARMM-GUI membrane-building tool was used to generate the systems and the simulation input files^54–56^. Each system consisted of the corresponding channel or transporter complex, a POPC lipid bilayer, TIP3 water molecules, and ions added to a concentration of 0.150 M of either NaCl or KCl (**Supp. Table 4**). The CHARMM36 m force field was used^57^, and all-atom MD simulations were run using GROMACS version 2022.2^58^. We performed minimization and equilibration (following the protocol outlined in CHARMM-GUI), including a 5000-step energy minimization, followed by 375 ps of MD equilibration at 1 fs/step, and then another 1500 ps of equilibration at 2 fs/step. After equilibration, a short 10 ns all-atom MD simulation was run (2 fs/step integration time) for the intact oligomer to generate slightly different monomer and oligomer configurations that served as initial conditions for each coarse-grained MD simulation (**Supp. Table 5-10**). We ran the production simulations in the NPT ensemble at 1 atmosphere and 310.15 K, using a C-rescale barostat with semi-isotropic pressure coupling and velocity rescaling for temperature coupling. For electrostatic interactions, we used Particle Mesh Ewald, and we applied a 12 Å cutoff for Lennard-Jones interactions with a switching function from 10-12 Å.

### Analysis of physicochemical properties of oligomeric interfaces

Snapshots of each system were obtained from the all-atom MD simulation as input for the analysis described in **Figure 1** and **Table 1**. Only the membrane-embedded regions were used, with residues defined as transmembrane by the corresponding UniProt entry in ExPASy ProtParam (**Supp. Table 1**). The oligomeric mass was determined by multiplying the protomer transmembrane region mass by the number of subunits that form the oligomer. The buried surface area, interfacial hydrogen bonds, and salt bridge residue pairs for each complex were calculated using the Protein Data Bank in Europe (PDBePISA) web server. A hydrogen bond is considered possible if the acceptor-hydrogen distance is less than 4 Å and the angle is between 90° and 270°. Salt bridges are considered possible for distances less than 4 Å between acidic (Asp, Glu) and basic (Arg, Lys, His) residues. Potential hydrogen bond and salt bridge residue pairs are reported with respect to chain A.

The surface complementarity (*S*_*C*_) coefficient for each oligomeric interface was calculated using the Python package Shape Complementarity and Available Surface Area (SCASA)^36^. Values of SC range from 0 to 1, with higher values indicating greater complementarity in surface shape.

### Coarse-grained molecular dynamics simulations

For the coarse-grained molecular dynamics (CGMD) simulations, systems were prepared using INSANE^60^ and run using the MARTINI 2.2 force field^37^ with GROMACS version 2020.2. Each system consisted of the protein (monomeric or oligomeric), a POPC lipid bilayer, water (10% antifreeze particles), and 150 mM ions. For each system, we prepared three independent runs for each state, using snapshots from the all-atom simulations to generate the initial systems. Details are provided in **Supp. Tables 5-10**. To maintain the protein’s secondary and tertiary structure, we applied an elastic network constraint with a force constant of 500 kJ/mol, lower and upper cutoffs of 0.5 and 0.9 nm, respectively, and an elastic bond decay factor of -ea 0 and decay power -ep 0.

Positional and orientational restraints were applied using PLUMED^62^ 2.6.2 to enable more efficient simulations, keep the system size smaller to reduce edge effects in membrane deformation analysis, and facilitate MOSAICS analysis (**Supp. Fig. 1**). The force constant was optimized to preserve natural tilting and wobbling in each system. To assess this, protein orientation was analyzed for simulations with and without restraints, using the *protein_orientation_mpi* tool in MOSAICS (**Supp. Fig. 2**). In this analysis, a vector parallel to the z-axis is obtained by defining parallel beads in the upper and lower regions of the protein. The beads are determined by first calculating the average coordinates of the protein across the simulations using the MOSAICS tool *mean_coords_mpi*. We then used the resulting PDB file with the average coordinates to identify bead pairs that define the vector. We obtained the bead pairs for this analysis using the MOSAICS tool *atoms_in_2_planes_mpi*, with a minimum z-axis distance between pairs of atoms of 2.5 nm and an XY cutoff distance of 0.3 nm. We selected the beads used for the analysis from chain A of Run 1 for each corresponding system. We then used the corresponding beads in each chain to determine the tilt and precession of each protomer. The same beads were used for the monomer state simulation. A full list of the bead restraints for each system is provided in **Supp. Table 11**.

To test for convergence of lipid sampling in each simulation, we calculated the degree of lipid exchange using the lipid mixing tool in MOSAICS (*lipid_mixing_mpi*). In this analysis, the XY position of each lipid is determined by using the geometric center of beads GL1 and GL2. We set a threshold distance of 1.1 nm from this center to define the lipids’ first shell. Across the trajectory, the lipid mixing analysis determines the number of lipids centered that had previously visited a lipid first shell and the average percentage of lipids that had previously occupied a lipid first shell. In general, 10 μs simulations in POPC result in 95% of lipids sampling every other lipid shell in the system (**Supp. Fig. 3**).

### Membrane thickness calculations

The 2D membrane thickness maps were calculated using the MOSAICS tool *membrane_thickness_mpi*. This analysis chooses a target leaflet and finds lipid pairs in the other leaflet that are closest to that lipid in the XY plane. The program returns the distance between the GL1 and GL2 geometric centers in the Z-direction. For the analysis, we used a grid area of 0.005 nm^2^, an XY distance cutoff of 0.5, a target atom radius of 0.26 nm, and a temperature of 310 K. This analysis yields the sample count and standard deviation per grid.

To determine whether a membrane grid element in a protein system was significantly perturbed compared to bulk, we first calculated the Z-score of the membrane thickness relative to a simulation of a bulk POPC lipid bilayer without any protein:

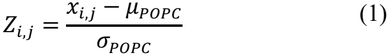

Where *μ*_*POPC*_ denotes the grand mean membrane thickness from a POPC-only simulation, and *σ*_*POPC*_ denotes the composite standard deviation, reflecting the variation in thickness across grid points and over the time course of the simulation (**Supp. Fig. 4**). In this form, the Z-score reports how many standard deviations a grid element is from a bulk POPC membrane thickness. Next, the fdrtool package in R was used to estimate the false discovery rate (FDR) corrected q-values for all grid points across all maps simultaneously using the calculated Z-score as input and the “fndr” cutoff method^63,64^. We selected a q-value of 0.05 as the cutoff for defining coordinates with membrane deformations. Using this, we calculated a Z-score cutoff of 0.058 to define significant deformation relative to bulk POPC, corresponding to an absolute membrane thickness difference of 0.014 nm or larger.

To calculate the distance that a grid element is from the protein surface, a 2D projection of the protein was estimated by calculating the convex hull of the undefined (NaN) values in the center of the MOSAICS maps using the concaveman package^65^. The distance of each coordinate in the MOSAICS map from the surface of the protein was calculated using the *st_distance* function in the sf package^66^.

### Rotating arc analysis to analyze different interfaces

To determine net changes in membrane deformation or lipid parameters across different protein faces, we developed a rotating arc analysis program in R using the 2D maps from the monomer simulations as input. Arcs with a defined radius (7 nm) and angular width (45° or 90°) were generated and rotated about each map’s center with a step size of 1°. Summary statistics of the parameters of interest are calculated for the grid points bounded by each arc, and the arc is assessed for overlap with the oligomeric interface, where the oligomeric interface residues are defined using the interfacial residues obtained via PDBePISA and later evaluated through 2D visual inspection (**Fig. 3A, B; Supp. Fig. 7**).

### Calculating membrane-dependent free energy changes

Using the MOSAICS tool, *membrane_thickness_mpi*, the free energy profile as a function of membrane thickness was calculated from the Boltzmann distribution of lipid-lipid distances in a POPC bilayer simulation:

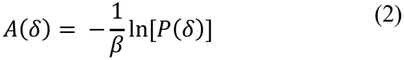

Where *A(δ)* is the free energy profile as a function of thickness, *P(δ)* is the probability of a lipid pair having a thickness *δ*, and *β* = 1/*k*_*B*_*T*, where *k*_*B*_ is the Boltzmann constant, and *T* is the temperature. The free energy profile is then shifted such that the minimum has a value of zero by using the equation:

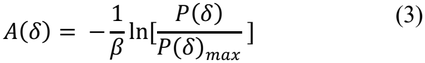

where *P(δ)*_*max*_ is the thickness with the highest probability (**Fig. 4E**). Applying this to the POPC-only simulation, we obtained the free energy cost of placing a pair of lipids at a specific thickness. Then, using the MOSAICS tool *bilayer_free_energ* on the membrane-protein maps, we determined the energy cost at each grid point based on the POPC-only free energy profile. The normalized lipid density used in the *bilayer_free_energ* analysis tool was determined using the *lipid_density_mpi* tool in MOSAICS.

### Lipid configurational analysis

We used the MOSAICS tools *lipid_orientation_mpi, interdigitation_mpi, lipid_distance_mpi*, and *lipid_contacts_mpi* to quantify lipid tilt, leaflet interdigitation, internal lipid distance, and lipid-water contacts, respectively, using an area per grid of 0.005 nm^2^ in each case^38^. For lipid orientation, we defined the vector between the lipid beads GL1/GL2 and C4A/C4B. To assess leaflet interdigitation, we used the C4A/C4B beads as the target leaflet and ranked the beads in the opposite leaflet. To measure the internal lipid distance, we used the distances between the atoms GL1-C4A and GL2-C4B. We measured lipid-water contacts between the water solvent and the hydrophobic region of the lipids (Beads C1A-C4A and C1B-C4B). MOSAICS generates a 2D map of each of these properties. To quantify changes relative to the bulk, we determined the mean value of each property in the bulk (defined for these analyses as grid points ≧ 4nm from the protein surfaces in each protein-POPC simulation) and subtracted it from each grid.

## ASSOCIATED CONTENT

### Supporting Information

The Supporting Information is available free of charge on the ACS Publications website:

Supplementary Figures (PDF) – Supplementary Figures 1-11

Supplementary Tables (PDF) – Supplementary Tables 1-11

## AUTHOR INFORMATION

### Author Contributions

The manuscript was written through contributions of all authors. All authors have given approval to the final version of the manuscript.

### Funding Sources

NIH R01GM120260

NIH R03NS133680

NIH T32HL007081

## ACKNOWLEDGMENT

We thank the Robertson Laboratory, Dr. Lucy Forrest, Dr. Giacomo Fiorin, Dr. Nathan Bernhardt, and the rest of the Forrest Laboratory for their insightful feedback during this project. This research was supported by the National Institutes of Health (NIH) through R01GM120260 (JLR & GDV), R03NS133680 (JLR), and the Principles of Cardiology Training Program (T32 HL007081 to GDV).

## ABBREVIATIONS

IMP: integral membrane protein
TM: transmembrane
POPC: palmityl-oleoyl phosphatidylcholine
MD: molecular dynamics
(CGMD): coarse-grained molecular dynamics

## Notes

### Competing Interest Statement

The authors have declared no competing interest.

## REFERENCES

(1) Singer, S. J.; Nicolson, G. L. The Fluid Mosaic Model of the Structure of Cell Membranes: Cell Membranes Are Viewed as Two-Dimensional Solutions of Oriented Globular Proteins and Lipids. Science 1972, 175 (4023), 720–731. 10.1126/science.175.4023.720

(2) Jost, P. C.; Griffith, O. H.; Capaldi, R. A.; Vanderkooi, G. Evidence for Boundary Lipid in Membranes. Proc. Natl. Acad. Sci. U. S. A. 1973, 70 (2), 480–484. 10.1073/pnas.70.2.480

(3) Marčelja, S. Lipid-Mediated Protein Interaction in Membranes. Biochim. Biophys. Acta BBA - Biomembr. 1976, 455 (1), 1–7. 10.1016/0005-2736(76)90149-8

(4) Israelachvili, J. N. Refinement of the Fluid-Mosaic Model of Membrane Structure. Biochim. Biophys. Acta BBA - Biomembr. 1977, 469 (2), 221–225. 10.1016/0005-2736(77)90185-7

(5) Lagüe, P.; Zuckermann, M. J.; Roux, B. Protein Inclusion in Lipid Membranes: A Theory Based on the Hypernetted Chain Integral Equation. Faraday Discuss. 1999, 111, 165–172. 10.1039/a807109h

(6) Lagüe, P.; Zuckermann, M. J.; Roux, B. Lipid-Mediated Interactions between Intrinsic Membrane Proteins: A Theoretical Study Based on Integral Equations. Biophys. J. 2000, 79 (6), 2867–2879. 10.1016/S0006-3495(00)76524-6

(7) Lagüe, P.; Zuckermann, M. J.; Roux, B. Lipid-Mediated Interactions between Intrinsic Membrane Proteins: Dependence on Protein Size and Lipid Composition. Biophys. J. 2001, 81 (1), 276–284. 10.1016/S0006-3495(01)75698-6

(8) Norimatsu, Y.; Hasegawa, K.; Shimizu, N.; Toyoshima, C. Protein–Phospholipid Interplay Revealed with Crystals of a Calcium Pump. Nature 2017, 545 (7653), 193–198. 10.1038/nature22357

(9) Feng, Z.; Alvarenga, O. E.; Accardi, A. Structural Basis of Closed Groove Scrambling by a TMEM16 Protein. Nat. Struct. Mol. Biol. 2024, 31 (10), 1468–1481. 10.1038/s41594-024-01284-9

(10) Guo, Y. R.; MacKinnon, R. Structure-Based Membrane Dome Mechanism for Piezo Mechanosensitivity. eLife 2017, 6, e33660. 10.7554/eLife.33660

(11) Lin, Y.-C.; Guo, Y. R.; Miyagi, A.; Levring, J.; MacKinnon, R.; Scheuring, S. Force-Induced Conformational Changes in PIEZO1. Nature 2019, 573 (7773), 230–234. 10.1038/s41586-019-1499-2

(12) Feng, Z.; Alvarenga, O. E.; Di Zanni, E.; Lee, S.; Khelashvili, G.; Accardi, A. Calcium Dependent Activation of the TMEM16F Scramblase and Ion Channel. Nat. Struct. Mol. Biol. 2026, 33 (4), 664–676. 10.1038/s41594-026-01789-5

(13) Blum, T. B.; Hahn, A.; Meier, T.; Davies, K. M.; Kühlbrandt, W. Dimers of Mitochondrial ATP Synthase Induce Membrane Curvature and Self-Assemble into Rows. Proc. Natl. Acad. Sci. 2019, 116 (10), 4250–4255. 10.1073/pnas.1816556116

(14) Davies, K. M.; Strauss, M.; Daum, B.; Kief, J. H.; Osiewacz, H. D.; Rycovska, A.; Zickermann, V.; Kühlbrandt, W. Macromolecular Organization of ATP Synthase and Complex I in Whole Mitochondria. Proc. Natl. Acad. Sci. 2011, 108 (34), 14121–14126. 10.1073/pnas.1103621108

(15) Davies, K. M.; Anselmi, C.; Wittig, I.; Faraldo-Gómez, J. D.; Kühlbrandt, W. Structure of the Yeast F1 Fo - ATP Synthase Dimer and Its Role in Shaping the Mitochondrial Cristae. Proc. Natl. Acad. Sci. 2012, 109 (34), 13602–13607. 10.1073/pnas.1204593109

(16) Anselmi, C.; Davies, K. M.; Faraldo-Gómez, J. D. --Mitochondrial ATP Synthase Dimers Spontaneously Associate Due to a Long-Range Membrane-Induced Force-. J. Gen. Physiol. 2018, 150 (5), 763–770. 10.1085/jgp.201812033

(17) Argudo, D.; Bethel, N. P.; Marcoline, F. V.; Wolgemuth, C. W.; Grabe, M. New Continuum Approaches for Determining Protein-Induced Membrane Deformations. Biophys. J. 2017, 112 (10), 2159–2172. 10.1016/j.bpj.2017.03.040

(18) Mondal, S.; Khelashvili, G.; Shan, J.; Andersen, O. S.; Weinstein, H. Quantitative Modeling of Membrane De-formations by Multihelical Membrane Proteins: Application to G-Protein Coupled Receptors. Biophys. J. 2011, 101 (9), 2092–2101. 10.1016/j.bpj.2011.09.037

(19) Zhou, Y. C.; Argudo, D.; Marcoline, F. V.; Grabe, M. A Computational Model of Protein Induced Membrane Morphology with Geodesic Curvature Driven Protein-Membrane Interface. J. Comput. Phys. 2020, 422, 109755. 10.1016/j.jcp.2020.109755

(20) Chadda, R.; Bernhardt, N.; Kelley, E. G.; Teixeira, S. C.; Griffith, K.; Gil-Ley, A.; Öztürk, T. N.; Hughes, L. E.; Forsythe, A.; Krishnamani, V.; Faraldo-Gómez, J. D.; Robertson, J. L. Membrane Transporter Dimerization Driven by Differential Lipid Solvation Energetics of Dissociated and Associated States. eLife 2021, 10, e63288. 10.7554/eLife.63288

(21) Ernst, M.; Orabi, E. A.; Stockbridge, R. B.; Faraldo-Gómez, J. D.; Robertson, J. L. Dimerization Mechanism of an Inverted-Topology Ion Channel in Membranes. Proc. Natl. Acad. Sci. 2023, 120 (47), e2308454120. 10.1073/pnas.2308454120

(22) Chadda, R.; Lee, T.; Mahoney-Kruszka, R.; Kelley, E. G.; Bernhardt, N.; Sandal, P.; Robertson, J. L. A Thermodynamic Analysis of CLC Transporter Dimerization in Lipid Bilayers. Proc. Natl. Acad. Sci. 2023, 120 (41), e2305100120. 10.1073/pnas.2305100120

(23) Bernhardt, N.; Ozturk, T. N.; Zhang, S.; Schwartz, N.; Chadda, R.; Gil-Ley, A.; Robertson, J. L.; Faraldo-Gómez, J. D. Molecular Basis for the Regulation of Membrane Proteins through Preferential Lipid Solvation. Nat. Chem. Biol. 2025. 10.1038/s41589-025-02032-w

(24) Coincon, M.; Uzdavinys, P.; Nji, E.; Dotson, D. L.; Winkelmann, I.; Abdul-Hussein, S.; Cameron, A. D.; Beckstein, O.; Drew, D. Crystal Structures Reveal the Molecular Basis of Ion Translocation in Sodium/Proton Antiporters. Nat. Struct. Mol. Biol. 2016, 23 (3), 248–255. 10.1038/nsmb.3164

(25) Dutzler, R.; Campbell, E. B.; MacKinnon, R. Gating the Selectivity Filter in ClC Chloride Channels. Science 2003, 300 (5616), 108–112. 10.1126/science.1082708

(26) Gonen, T.; Cheng, Y.; Sliz, P.; Hiroaki, Y.; Fujiyoshi, Y.; Harrison, S. C.; Walz, T. Lipid–Protein Interactions in Double-Layered Two-Dimensional AQP0 Crystals. Nature 2005, 438 (7068), 633–638. 10.1038/nature04321

(27) Gotfryd, K.; Boesen, T.; Mortensen, J. S.; Khelashvili, G.; Quick, M.; Terry, D. S.; Missel, J. W.; LeVine, M. V.; Gourdon, P.; Blanchard, S. C.; Javitch, J. A.; Weinstein, H.; Loland, C. J.; Nissen, P.; Gether, U. X-Ray Structure of LeuT in an Inward-Facing Occluded Conformation Reveals Mechanism of Substrate Release. Nat. Commun. 2020, 11 (1), 1005. 10.1038/s41467-020-14735-w

(28) Ilgü, H.; Jeckelmann, J.-M.; Kalbermatter, D.; Ucurum, Z.; Lemmin, T.; Fotiadis, D. High-Resolution Structure of the Amino Acid Transporter AdiC Reveals Insights into the Role of Water Molecules and Networks in Oligomerization and Substrate Binding. BMC Biol. 2021, 19 (1), 179. 10.1186/s12915-021-01102-4

(29) Lee, Y.; Nishizawa, T.; Yamashita, K.; Ishitani, R.; Nureki, O. Structural Basis for the Facilitative Diffusion Mechanism by SemiSWEET Transporter. Nat. Commun. 2015, 6 (1), 6112. 10.1038/ncomms7112

(30) Ma, D.; Lu, P.; Yan, C.; Fan, C.; Yin, P.; Wang, J.; Shi, Y. Structure and Mechanism of a Glutamate–GABA Antiporter. Nature 2012, 483 (7391), 632–636. 10.1038/nature10917

(31) Stockbridge, R. B.; Kolmakova-Partensky, L.; Shane, T.; Koide, A.; Koide, S.; Miller, C.; Newstead, S. Crystal Structures of a Double-Barrelled Fluoride Ion Channel. Nature 2015, 525 (7570), 548–551. 10.1038/nature14981

(32) Wang, Y.; Huang, Y.; Wang, J.; Cheng, C.; Huang, W.; Lu, P.; Xu, Y.-N.; Wang, P.; Yan, N.; Shi, Y. Structure of the Formate Transporter FocA Reveals a Pentameric Aquaporin-like Channel. Nature 2009, 462 (7272), 467–472. 10.1038/nature08610

(33) Zhou, Y.; Morais-Cabral, J. H.; Kaufman, A.; MacKinnon, R. Chemistry of Ion Coordination and Hydration Revealed by a K+ Channel–Fab Complex at 2.0 Å Resolution. Nature 2001, 414 (6859), 43–48. 10.1038/35102009

(34) Edwards, N.; Anderson, C. M. H.; Conlon, N. J.; Watson, A. K.; Hall, R. J.; Cheek, T. R.; Embley, T. M.; Thwaites, D. T. Resculpting the Binding Pocket of APC Superfamily LeuT-Fold Amino Acid Transporters. Cell. Mol. Life Sci. 2018, 75 (5), 921–938. 10.1007/s00018-017-2677-8

(35) Krishnamurthy, H.; Gouaux, E. X-Ray Structures of LeuT in Substrate-Free Outward-Open and Apo In-ward-Open States. Nature 2012, 481 (7382), 469–474. 10.1038/nature10737

(36) Lawrence, M. C.; Colman, P. M. Shape Complementarity at Protein/Protein Interfaces. J. Mol. Biol. 1993, 234 (4), 946–950. 10.1006/jmbi.1993.1648

(37) De Jong, D. H.; Singh, G.; Bennett, W. F. D.; Arnarez, C.; Wassenaar, T. A.; Schäfer, L. V.; Periole, X.; Tieleman, D. P.; Marrink, S. J. Improved Parameters for the Martini Coarse-Grained Protein Force Field. J. Chem. Theory Comput. 2013, 9 (1), 687–697. 10.1021/ct300646g

(38) Bernhardt, N.; Faraldo-Gómez, J. D. MOSAICS: A Software Suite for Analysis of Membrane Structure and Dynamics in Simulated Trajectories. Biophys. J. 2023, 122 (11), 2023–2040. 10.1016/j.bpj.2022.11.005

(39) Benjamini, Y.; Hochberg, Y. Controlling the False Discovery Rate: A Practical and Powerful Approach to Multiple Testing. J. R. Stat. Soc. Ser. B Stat. Methodol. 1995, 57 (1), 289–300. 10.1111/j.2517-6161.1995.tb02031.x

(40) Efron, B. Large-Scale Simultaneous Hypothesis Testing: The Choice of a Null Hypothesis. J. Am. Stat. Assoc. 2004, 99 (465), 96–104. 10.1198/016214504000000089

(41) Strimmer, K. Fdrtool: A Versatile R Package for Estimating Local and Tail Area-Based False Discovery Rates. Bioinformatics 2008, 24 (12), 1461–1462. 10.1093/bioinformatics/btn209

(42) Strimmer, K. A Unified Approach to False Discovery Rate Estimation. BMC Bioinformatics 2008, 9 (1), 303. 10.1186/1471-2105-9-303

(43) Andersen, O. S.; Koeppe, R. E. Bilayer Thickness and Membrane Protein Function: An Energetic Perspective. Annu. Rev. Biophys. Biomol. Struct. 2007, 36 (1), 107–130. 10.1146/annurev.biophys.36.040306.132643

(44) Hite, R. K.; Chiu, P.-L.; Schuller, J. M.; Walz, T. Effect of Lipid Head Groups on Double-Layered Two-Dimensional Crystals Formed by Aquaporin-0. PLOS ONE 2015, 10 (1), e0117371. 10.1371/journal.pone.0117371

(45) Scheuring, S.; Buzhynskyy, N.; Jaroslawski, S.; Gonçalves, R. P.; Hite, R. K.; Walz, T. Structural Models of the Supramolecular Organization of AQP0 and Connexons in Junctional Microdomains. J. Struct. Biol. 2007, 160 (3), 385–394. 10.1016/j.jsb.2007.07.009

(46) Guo, Y. Be Cautious with Crystal Structures of Membrane Proteins or Complexes Prepared in Detergents. Crystals 2020, 10 (2), 86. 10.3390/cryst10020086

(47) Perez, C.; Khafizov, K.; Forrest, L. R.; Krämer, R.; Ziegler, C. The Role of Trimerization in the Osmoregulated Betaine Transporter BetP. EMBO Rep. 2011, 12 (8), 804–810. 10.1038/embor.2011.102

(48) Coleman, J. A.; Green, E. M.; Gouaux, E. X-Ray Structures and Mechanism of the Human Serotonin Transporter. Nature 2016, 532 (7599), 334–339. 10.1038/nature17629

(49) Li, Y.; Wang, X.; Meng, Y.; Hu, T.; Zhao, J.; Li, R.; Bai, Q.; Yuan, P.; Han, J.; Hao, K.; Wei, Y.; Qiu, Y.; Li, N.; Zhao, Y. Dopamine Reuptake and Inhibitory Mechanisms in Human Dopamine Transporter. Nature 2024, 632 (8025), 686–694. 10.1038/s41586-024-07796-0

(50) Paul, A.; Shukla, D. Oligomerization of Monoamine Transporters. In Macromolecular Protein Complexes V; Harris, J. R., Marles-Wright, J., Eds.; Subcellular Biochemistry; Springer International Publishing: Cham, 2024; Vol. 104, pp 119–137. 10.1007/978-3-031-58843-3_7

(51) Rudin, D.; Luethi, D.; Niello, M.; Yang, J.-W.; Burger, I.; Sandtner, W.; Birner-Gruenberger, R.; Schütz, G. J.; Sitte, H. H. Cell Membrane Cholesterol Affects Serotonin Transporter Efflux Due to Altered Transporter Oligomerization. Mol. Psychiatry 2026, 31 (2), 963–975. 10.1038/s41380-025-03201-y

(52) Zhang, H.; Yin, Y.-L.; Dai, A.; Zhang, T.; Zhang, C.; Wu, C.; Hu, W.; He, X.; Pan, B.; Jin, S.; Yuan, Q.; Wang, M.-W.; Yang, D.; Xu, H. E.; Jiang, Y. Dimerization and Antidepressant Recognition at Noradrenaline Transporter. Nature 2024, 630 (8015), 247–254. 10.1038/s41586-024-07437-6

(53) Webb, B.; Sali, A. Comparative Protein Structure Modeling Using MODELLER. Curr. Protoc. Bioinforma. 2016, 54 (1). 10.1002/cpbi.3

(54) Jo, S.; Kim, T.; Iyer, V. G.; Im, W. CHARMM-GUI: A Web-based Graphical User Interface for CHARMM. J. Comput. Chem. 2008, 29 (11), 1859–1865. 10.1002/jcc.20945

(55) Lee, J.; Cheng, X.; Swails, J. M.; Yeom, M. S.; Eastman, P. K.; Lemkul, J. A.; Wei, S.; Buckner, J.; Jeong, J. C.; Qi, Y.; Jo, S.; Pande, V. S.; Case, D. A.; Brooks, C. L.; MacKerell, A. D.; Klauda, J. B.; Im, W. CHARMM-GUI Input Generator for NAMD, GROMACS, AMBER, OpenMM, and CHARMM/OpenMM Simulations Using the CHARMM36 Additive Force Field. J. Chem. Theory Comput. 2016, 12 (1), 405–413. 10.1021/acs.jctc.5b00935

(56) Wu, E. L.; Cheng, X.; Jo, S.; Rui, H.; Song, K. C.; Dávila-Contreras, E. M.; Qi, Y.; Lee, J.; Monje-Galvan, V.; Venable, R. M.; Klauda, J. B.; Im, W. CHARMM-GUI Membrane Builder toward Realistic Biological Membrane Simulations. J. Comput. Chem. 2014, 35 (27), 1997–2004. 10.1002/jcc.23702

(57) Huang, J.; Rauscher, S.; Nawrocki, G.; Ran, T.; Feig, M.; De Groot, B. L.; Grubmüller, H.; MacKerell, A. D. CHARMM36m: An Improved Force Field for Folded and Intrinsically Disordered Proteins. Nat. Methods 2017, 14 (1), 71–73. 10.1038/nmeth.4067

(58) Abraham, M. J.; Murtola, T.; Schulz, R.; Páll, S.; Smith, J. C.; Hess, B.; Lindahl, E. GROMACS: High Performance Molecular Simulations through Multi-Level Parallelism from Laptops to Supercomputers. SoftwareX 2015, 1–2, 19–25. 10.1016/j.softx.2015.06.001

(59) Krissinel, E.; Henrick, K. Inference of Macromolecular Assemblies from Crystalline State. J. Mol. Biol. 2007, 372 (3), 774–797. 10.1016/j.jmb.2007.05.022

(60) Wassenaar, T. A.; Ingólfsson, H. I.; Böckmann, R. A.; Tieleman, D. P.; Marrink, S. J. Computational Lipidomics with Insane : A Versatile Tool for Generating Custom Membranes for Molecular Simulations. J. Chem. Theory Comput. 2015, 11 (5), 2144–2155. 10.1021/acs.jctc.5b00209

(61) Marrink, S. J.; Risselada, H. J.; Yefimov, S.; Tieleman, D. P.; De Vries, A. H. The MARTINI Force Field: Coarse Grained Model for Biomolecular Simulations. J. Phys. Chem. B 2007, 111 (27), 7812–7824. 10.1021/jp071097f

(62) Bonomi, M.; Branduardi, D.; Bussi, G.; Camilloni, C.; Provasi, D.; Raiteri, P.; Donadio, D.; Marinelli, F.; Pietrucci, F.; Broglia, R. A.; Parrinello, M. PLUMED: A Portable Plugin for Free-Energy Calculations with Molecular Dynamics. Comput. Phys. Commun. 2009, 180 (10), 1961–1972. 10.1016/j.cpc.2009.05.011

(63) R Core Team. R: A Language and Environment for Statistical Computing; R Foundation for Statistical Computing: Vienna, Austria, 2025.

(64) Strimmer, K. Fdrtool: A Versatile R Package for Estimating Local and Tail Area-Based False Discovery Rates. Bioinformatics 2008, 24 (12), 1461–1462. 10.1093/bioinformatics/btn209

(65) Gombin, J.; Vaidyanathan, R.; Agafonkin, V. Concaveman: A Very Fast 2D Concave Hull Algorithm, 2017, 1.2.0. 10.32614/CRAN.package.concaveman

(66) Pebesma, E. Simple Features for R: Standardized Support for Spatial Vector Data. R J. 2018, 10 (1), 439. 10.32614/RJ-2018-009 1.2.0. https://doi.org/10.32614/CRAN.package.concaveman

(66) Pebesma, E. Simple Features for R: Standardized Support for Spatial Vector Data. R J. 2018, 10 (1), 439. 10.32614/RJ-2018-009

