## Supplementary Information for "Structural deformations of lipid bilayers around membrane proteins inform oligomerization type"

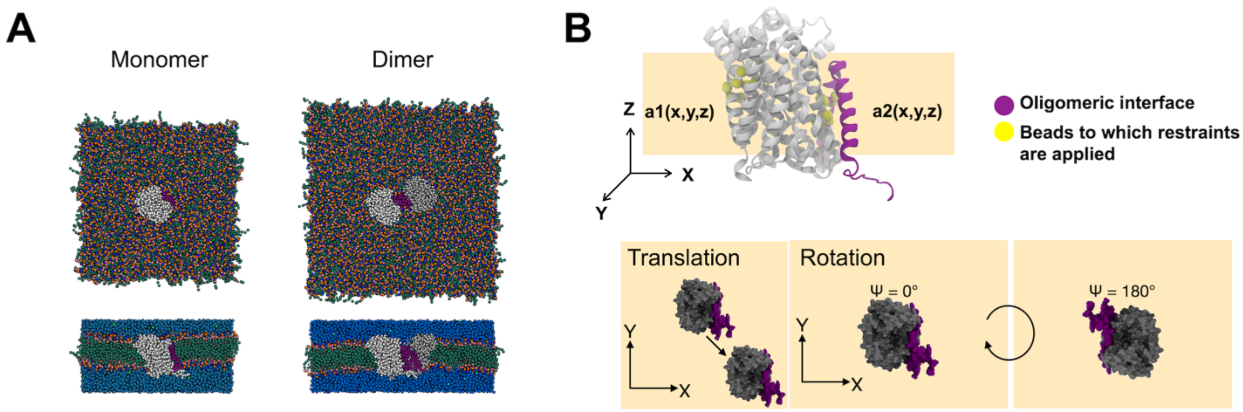


**Supplementary Figure 1. Coarse-grained molecular dynamics simulations with positional and orientational restraints.** (A) Top and side views of a representative system for the molecular dynamics simulations. Protein atoms are colored white and gray, with the oligomeric interface highlighted in purple. Lipid hydrophobic beads are colored green, headgroups are blue and orange, and water is blue. (B) To optimize simulation and analysis efficiency, positional and orientational restraints were applied to specific protein beads (yellow), restricting their x, y, and z positions and limiting protein rotation, as well as lateral and vertical movement. System specifications are listed in Supp. Tables 5-10, and bead restraints are listed in Supp. Table 11.

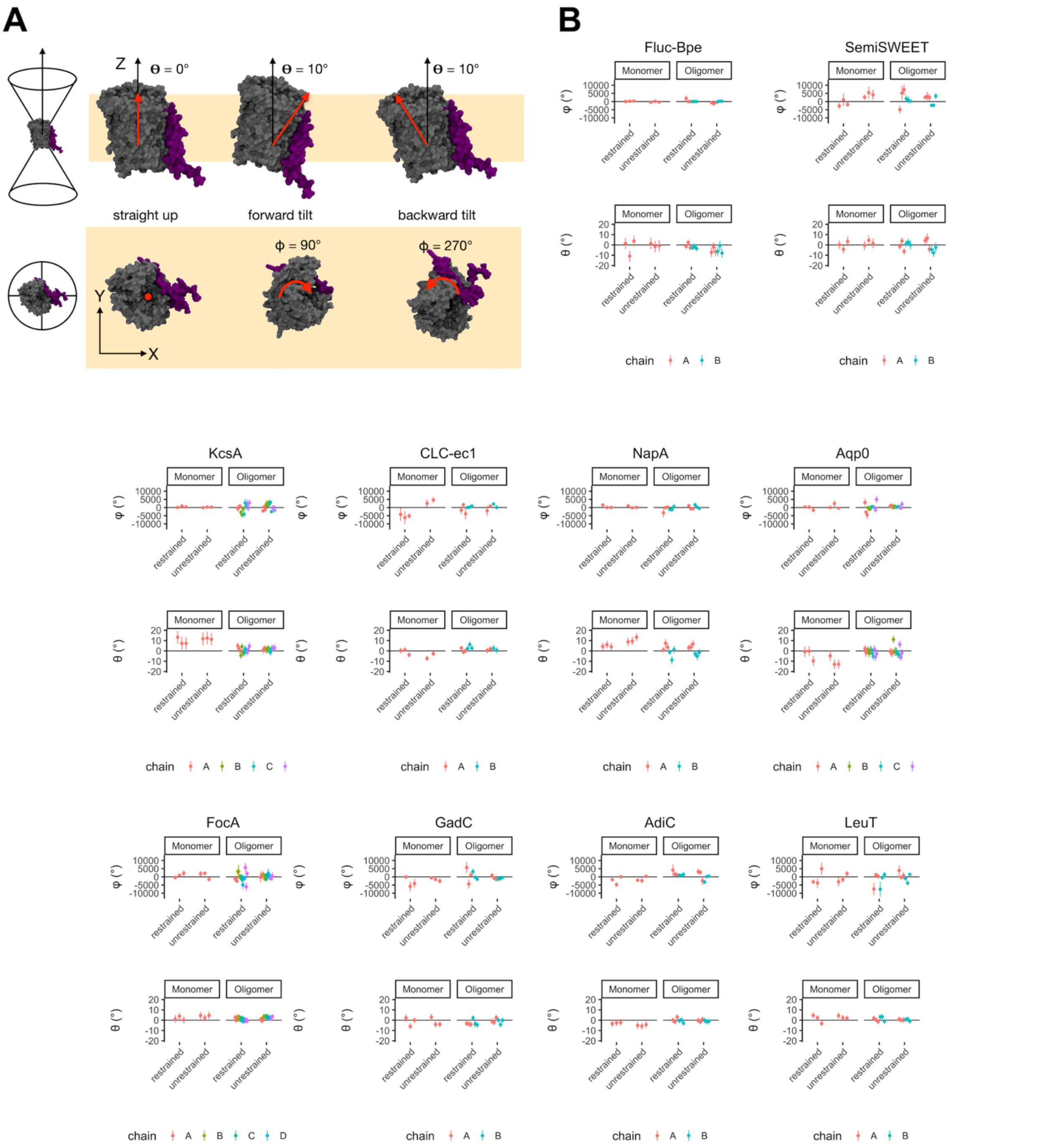


**Supplementary Figure 2. Analysis of protein tilt and precession with and without restraints.** To ensure the positional and orientational restraints applied to the system did not limit conformational sampling, we analyzed protein orientation in the restrained and unrestrained systems. (A) Schematic showing tilt (θ) and precession (ϕ) angles. (B) Comparison of the net tilt and precession for each protein chain in restrained and unrestrained simulations. A summary of final tilt angles is listed in Supp. Tables 2 & 3.

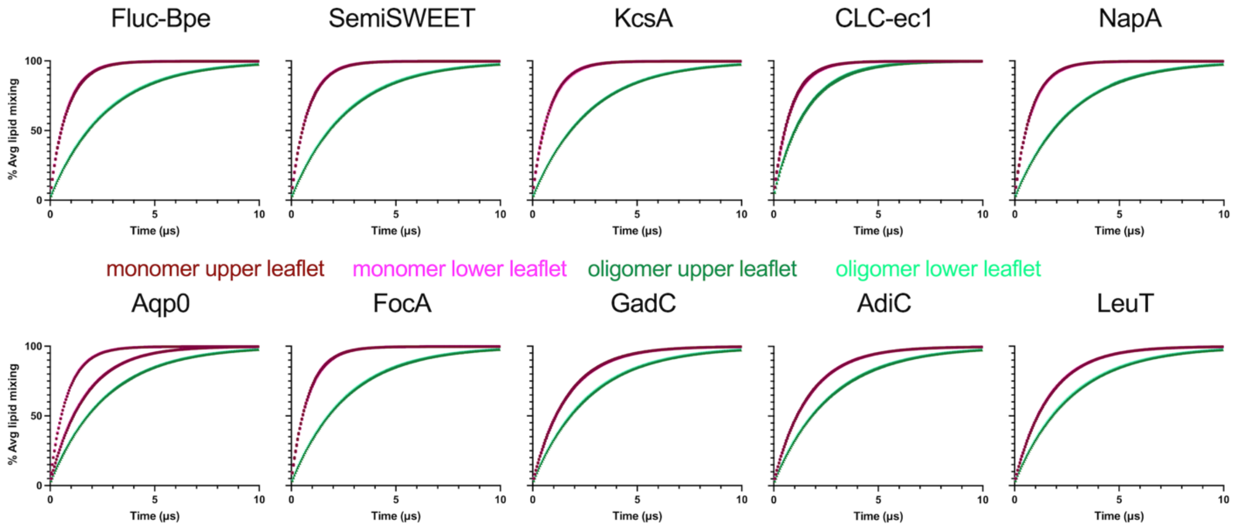


**Supplementary Figure 3. Convergence of lipid mixing.** MOSAICS calculation of lipid mixing in each simulation, separated into upper- and lower-leaflet analysis for monomer and oligomer simulations. For Aqp0, two monomer curves are observed because the simulation replicates used different system sizes.

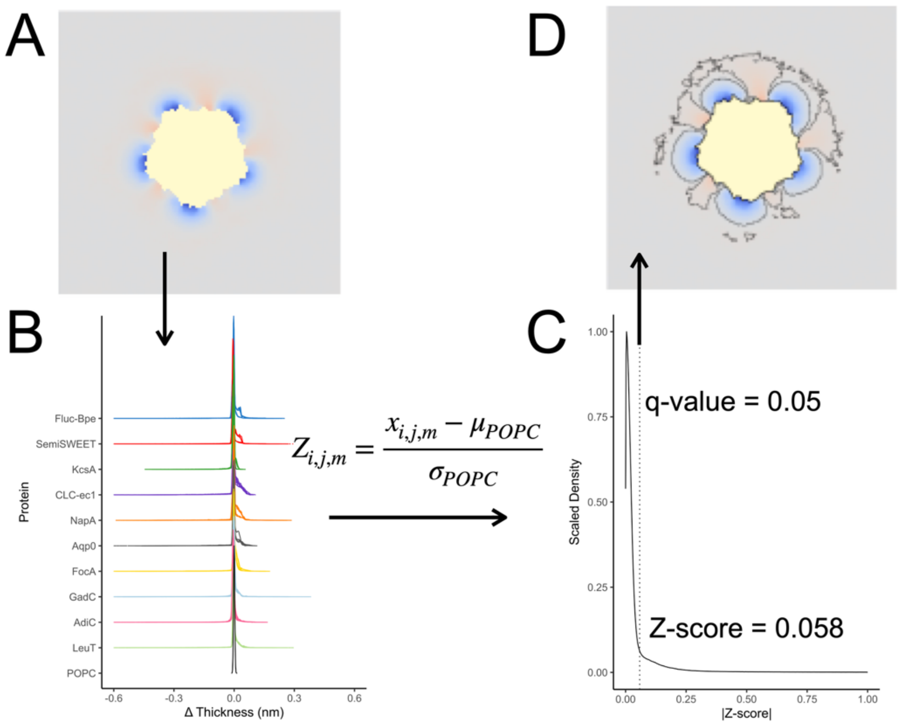


**Supplementary Figure 4. Identifying statistically significant membrane defects using Z-score statistics and false discovery rate (FDR).** (A) An example of a 2D membrane thickness map for one simulation. (B) Probability density distributions of membrane thickness deviations over all grid points ($i$, $j$) and for each map $m$. The Z-score describes how many standard deviations a value is from a reference distribution. Here, the reference distribution is the POPC bilayer alone, providing the mean and composite standard deviation of membrane thickness. (C) Scaled density plot of the combined set of Z-scores used as input for FDR estimation. The Z-score threshold for q-values ≤ 0.05 is shown by the dotted line, corresponding to a Z-score = 0.058. (D) Membrane thickness map in (A) with statistically significant defects outlined.

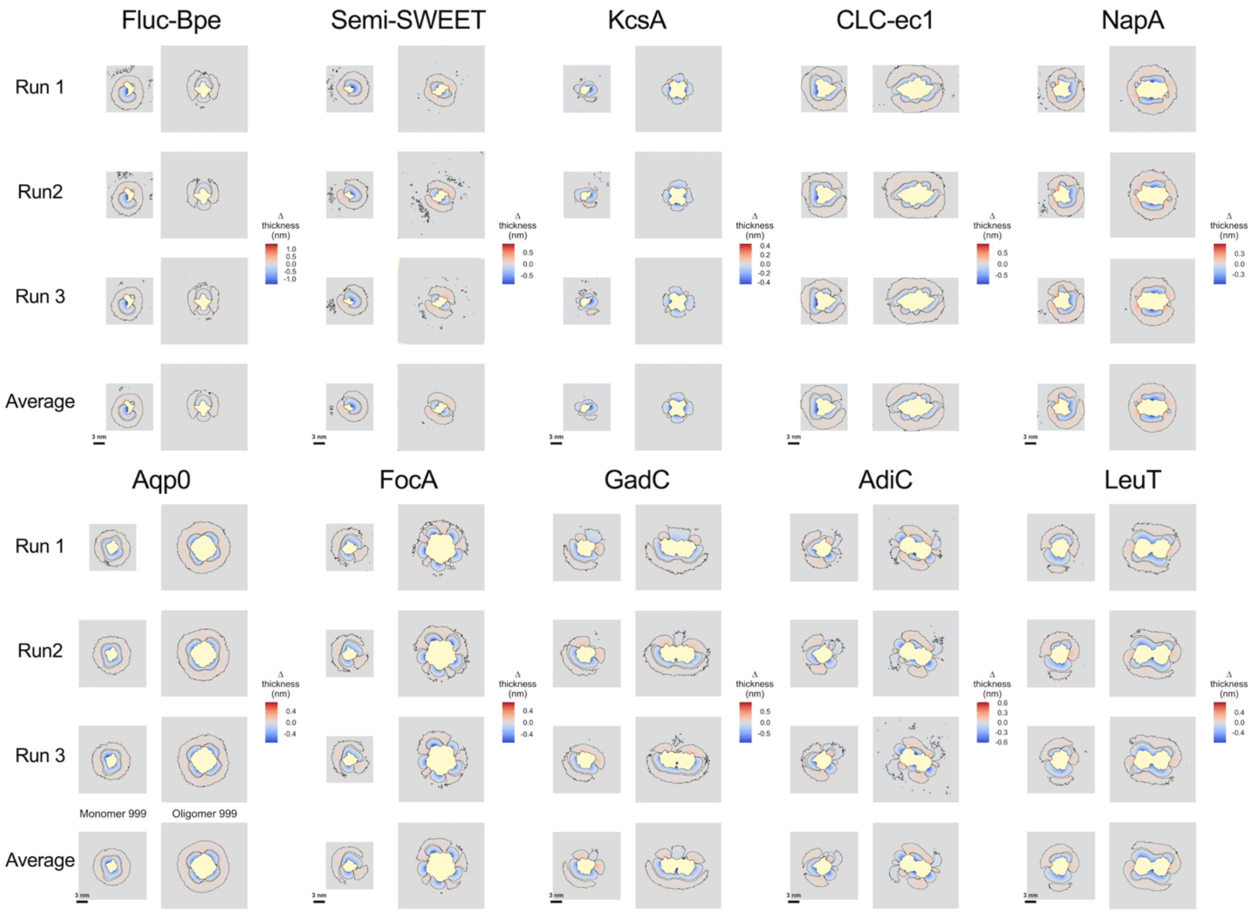


**Supplementary Figure 5. Membrane thickness maps highlighting significant defects.** 2D membrane thickness maps for each channel and transporter under study with defined deformed regions surrounded by black lines for runs 1, 2, 3, and the average of the three runs.

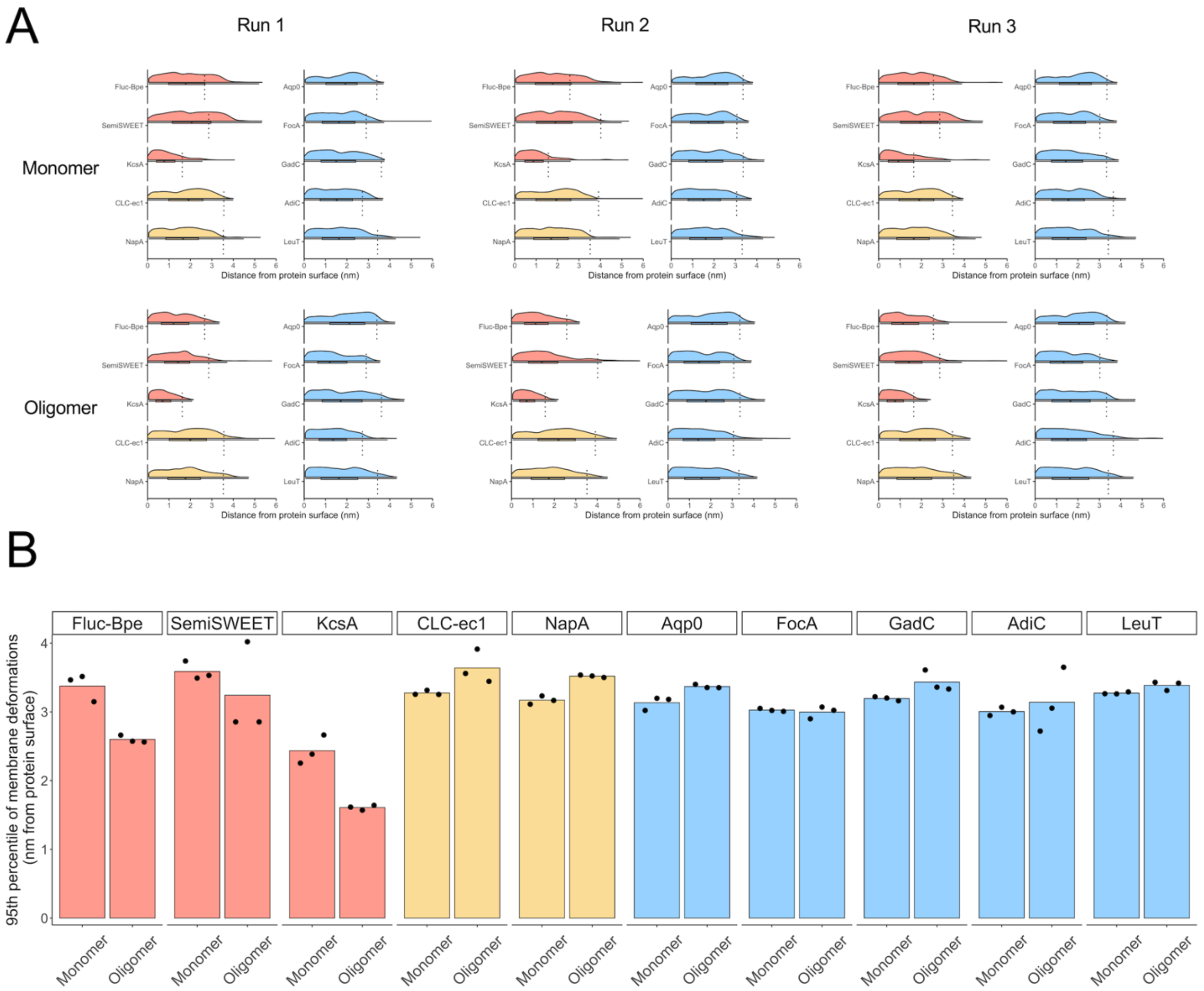


**Supplementary Figure 6. Analysis of the extent of membrane defects around each protein.** (A) Probability density distributions of the distance of significantly perturbed grid points from the surface of the protein, for each independent simulation. (B) 95$^{th}$ percentile of the distance of significantly perturbed grid points from the surface of each protein.

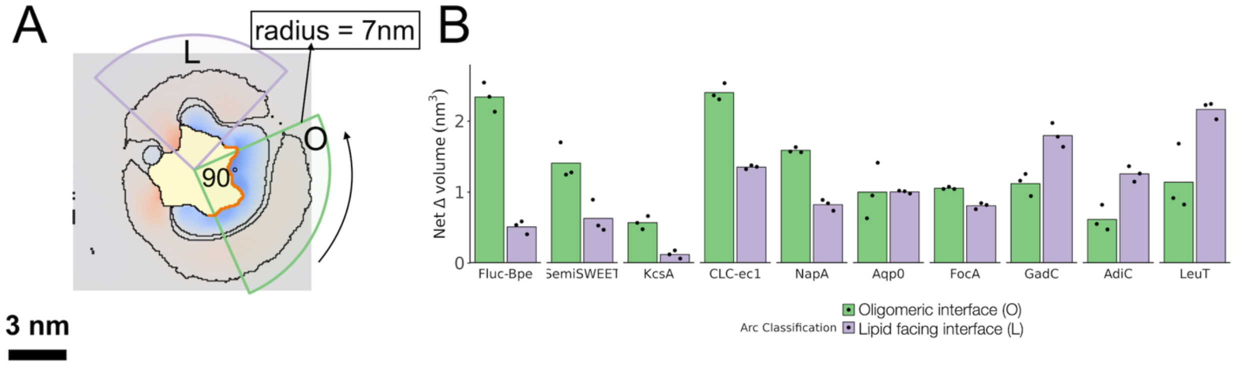


**Supplementary Figure 7. Arc analysis around monomer interfaces.** (A) Depiction of the rotating arc analysis method. An arc is defined with a specific radius and angle and rotated around the center of the protein to calculate the net deformation in each area. The arcs bounding the maximum deformed volume are shown for the oligomeric interface (O) and lipid-facing face (L). (B) Quantification of the arc analysis for a 90˚ wedge with a 7 nm radius for the O-L data presented in Fig. 3B.

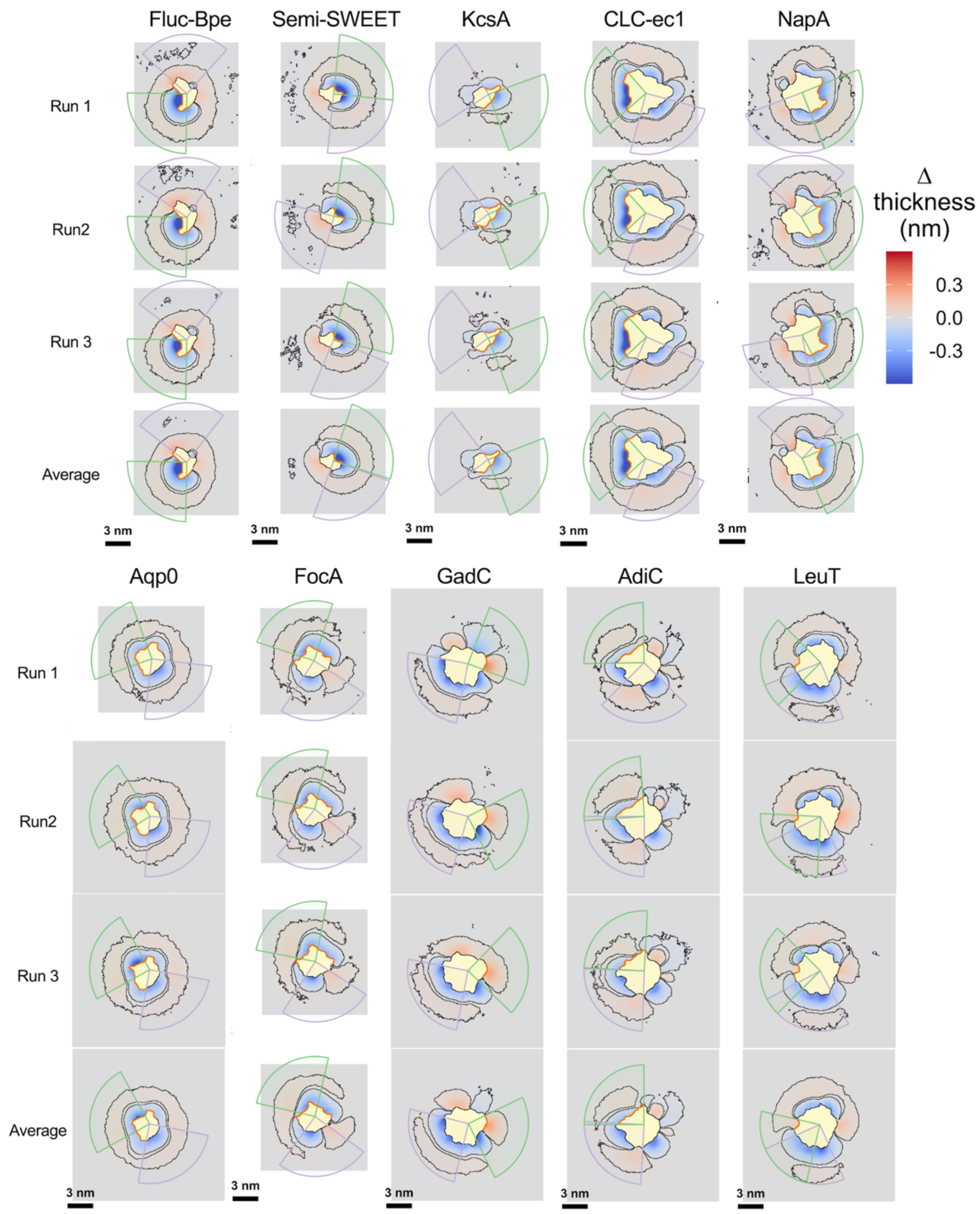


**Supplementary Figure 8. Monomer membrane thickness maps with maximum defects at oligomeric and lipid-facing interfaces.** Membrane thickness maps for 3 replicates and the average, with arcs identifying the oligomerization interface (green) and the lipid-exposed interface (purple) that exhibit maximum defects; arc size 90˚ and radius 7 nm. The orange line highlights the oligomerization interface.

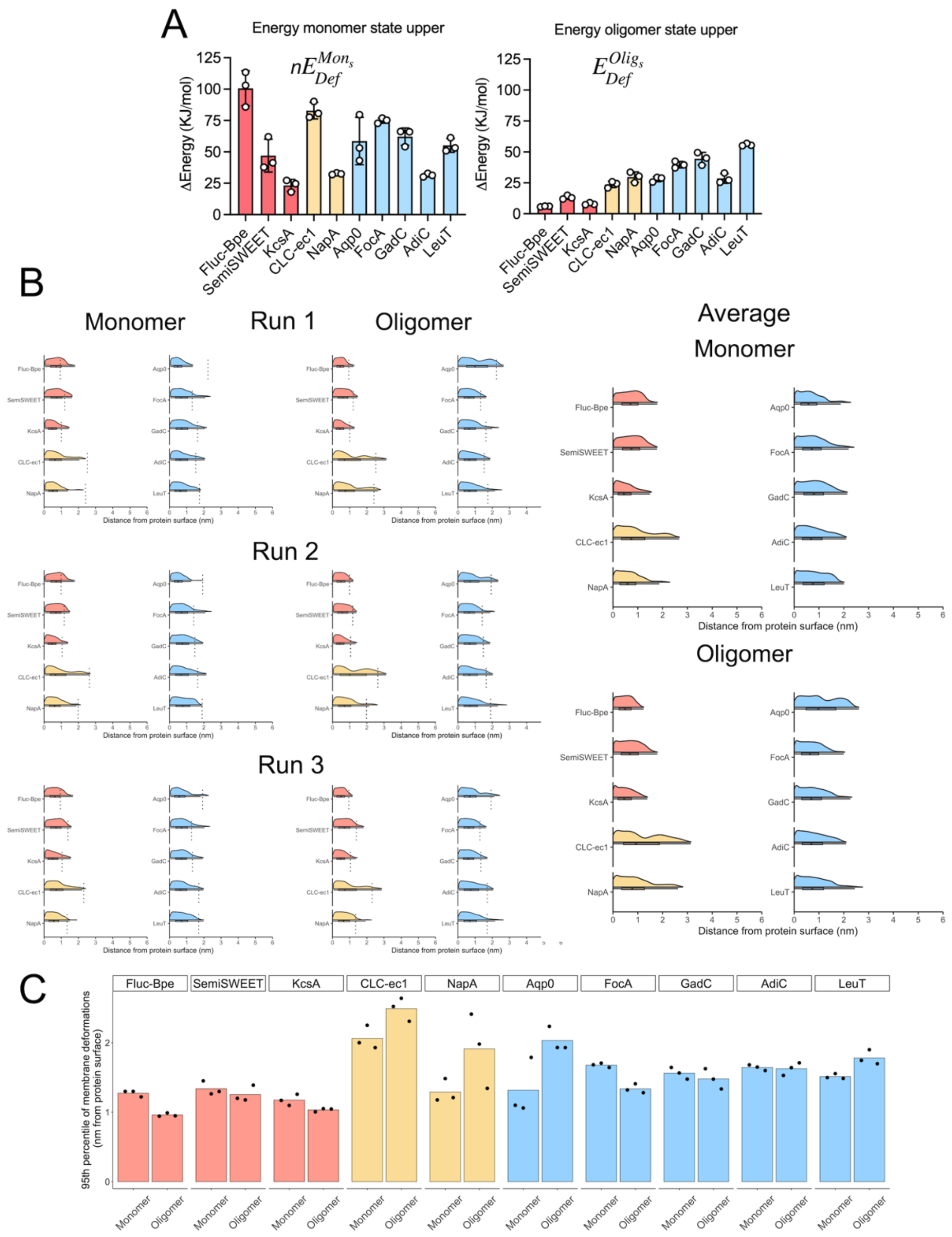


**Supplementary Figure 9. Energetic contributions of membrane defects.** (A) Energetic contribution of membrane deformations around n*monomers and the oligomeric states. (B) Probability density distributions of grid points with a non-zero energetic penalty as a function of the distance from the protein surface. Results are shown for each independent run and the average of both monomer and oligomer simulations. (C) 95th percentile distances of grid points with a non-zero energetic penalty from the protein surface.

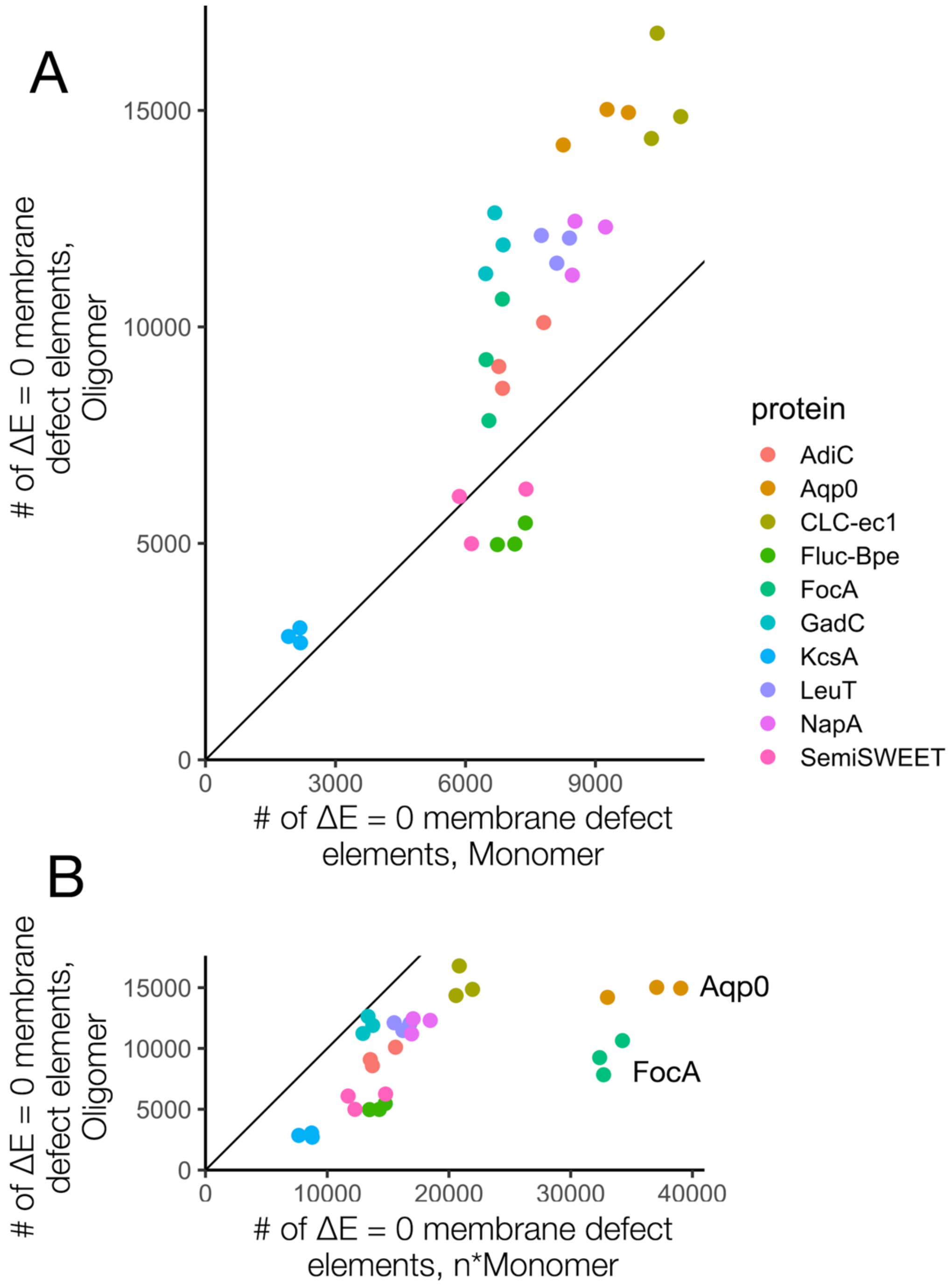


**Supplementary Figure 10. Correlation plots of grid points with statistically significant changes in membrane thickness and no energetic penalty.** (A) Comparison of Oligomer vs. Monomer. (B) Comparison of Oligomer vs. n*Monomer. The diagonal line represents Y = X. Each dot represents one pair of simulations.

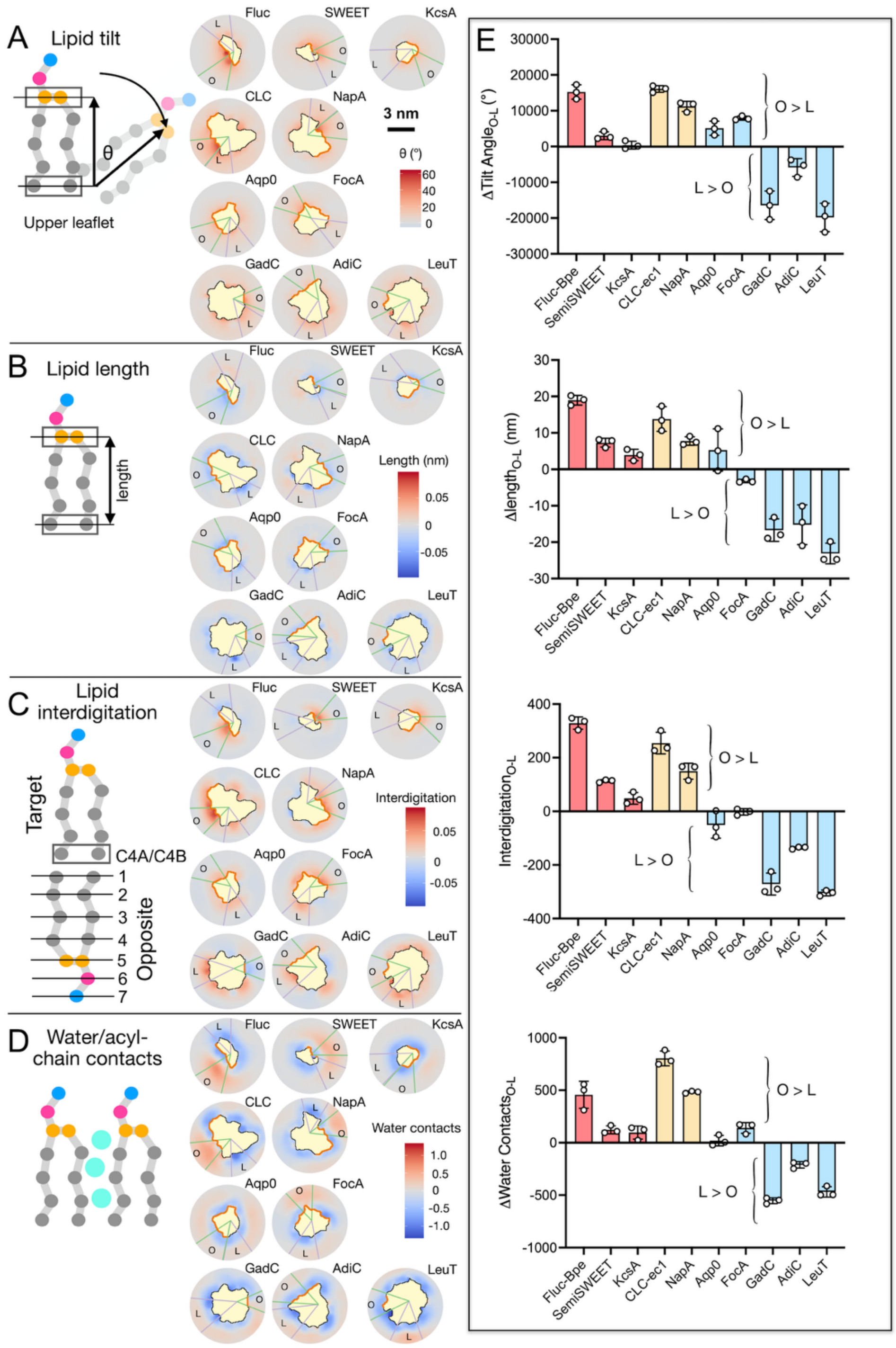


**Supplementary Figure 11. Analysis of lipid configurations.** Average lipid configurations were analyzed using MOSAICS. (A) Lipid tilt angle, defined as the angle between the membrane normal and a vector defined by the selected beads (boxed). (B) Lipid length, defined as the distance between the selected beads (boxed). (C) Lipid interdigitation is measured by defining beads C4A/C4B in the target leaflet and measuring their contact with the ranked beads on the opposite leaflet. (D) Lipid-water contact is measured by counting the number of water molecules that enter in contact with the hydrophobic region of the lipids (gray beads). For (A-D), maps corresponding to the upper leaflet results are presented with 45˚ arcs. (E) Rotating arc analysis for each lipid configuration parameter, showing the maximum defect at the oligomeric interface (O) minus the defect at the lipid-facing face (L), i.e., O-L.

### SUPPLEMENTARY TABLES:

### Structural deformations of lipid bilayers around membrane proteins inform oligomerization type

Gladys Díaz Vázquez[1], Samson G. F. Condon[2], Janice L. Robertson[1]*

[1]Department of Biochemistry and Molecular Biophysics, Washington University School of Medicine, St. Louis, MO 63110, [2]Independent Scholar

**Supplementary Table 1.** Protein systems used in this study, listing the membrane-embedded residues included in Figure 1 and Table 1.

| Protein | Uniprot ID | Membrane embedded residues |
|---|---|---|
| Fluc-Bpe | Q7VYU0 | 7-29, 36-57, 65-94, 98-126 |
| SemiSWEET | P0DMV3 | 4-27, 35-55, 60-82 |
| KcsA | P0A334 | 28-50, 62-72, 73-80, 88-111 |
| CLC-ec1 | P37019 | 33-69, 77-100, 109-116, 124-141, 148-166, 177-189, 193-201, 215-232, 253-281, 288-309, 330-349, 355-376, 387-401, 402-404, 405-416, 417-421, 422-438 |
| NapA | Q5SIA2 | 6-24, 31-49, 55-75, 87-105, 117-136, 148-168, 180-203, 215-248, 260-278, 290-314, 353-375 |
| Aqp0 | Q6J8I9 | 9-32, 39-61, 62-67, 68-78, 85-107, 127-147, 160-176, 177-183, 184-194, 201-219 |
| FocA | P0AC25 | 31-56, 65-85, 113-135, 161-181, 188-205, 250-276 |
| GadC | P63235 | 14-36, 41-64, 86-112, 127-147, 152-180, 202-225, 230-259, 289-322, 338-359, 363-396, 410-430, 444-467 |
| AdiC | P60061 | 12-24, 41-62, 84-113, 124-142, 146-171, 189-204, 223-247, 273-301, 325-343, 347-370, 385-404, 409-427 |
| LeuT | O67854 | 12-29, 41-61, 92-111, 166-185, 197-215, 243-268, 289-314, 342-367, 379-399, 405-427, 447-469, 481-504 |

**Supplementary Table 2.** Monomer tilt angles with respect to the membrane perpendicular z-axis.

| | Tilt angle (º) | | | |
|---|---|---|---|---|
| Protein | Run 1 | Run 2 | Run 3 | Mean ± standard deviation |
| Fluc-Bpe | 17.48 | 17.42 | 16.94 | 17.28 ± 0.29 |
| SemiSWEET | 8.41 | 8.42 | 8.83 | 8.56 ± 0.24 |
| KcsA | 17.79 | 16.40 | 18.79 | 17.66 ± 1.20 |
| CLC-ec1 | 5.09 | 4.74 | 4.58 | 4.80 ± 0.26 |
| NapA | 10.66 | 13.12 | 13.39 | 12.39 ± 1.50 |
| Aqp0 | 18.61 | 9.13 | 9.60 | 12.45 ± 5.34 |
| FocA | 7.72 | 7.32 | 7.17 | 7.40 ± 0.28 |
| GadC | 10.07 | 6.57 | 5.25 | 7.30 ± 2.49 |
| AdiC | 8.03 | 6.85 | 7.85 | 7.58 ± 0.64 |
| LeuT | 5.75 | 4.89 | 5.06 | 5.24 ± 0.45 |

**Supplementary Table 3.** Difference in tilt angles between monomer and oligomer (chain A).

| | Tilt angle (º) | | | |
|---|---|---|---|---|
| Protein | Run 1 | Run 2 | Run 3 | Mean ± standard deviation |
| Fluc-Bpe | 12.54 | 9.11 | 8.30 | 9.98 ± 2.25 |
| SemiSWEET | 1.94 | 1.82 | 2.64 | 2.13 ± 0.44 |
| KcsA | 11.25 | 9.88 | 13.12 | 11.42 ± 1.63 |
| CLC-ec1 | 1.06 | 0.10 | 0.01 | 0.39 ± 0.58 |
| NapA | 5.03 | 0.98 | 4.70 | 3.57 ± 2.25 |
| Aqp0 | 12.03 | 0.58 | 3.06 | 5.22 ± 6.02 |
| FocA | 3.43 | 2.64 | 2.16 | 2.74 ± 0.64 |
| GadC | 5.38 | 1.68 | -0.30 | 2.25 ± 2.88 |
| AdiC | 2.98 | 2.20 | 2.70 | 2.62 ± 0.40 |
| LeuT | 1.46 | 0.86 | 0.60 | 0.97 ± 0.44 |

**Supplementary Table 4.** System details for all-atom oligomer molecular dynamics simulations.

| **Components** | **Fluc-Bpe** | **Semi SWEET** | **KcsA** | **CLC-ec1** | **NapA** | **Aqp0** | **FocA** | **GadC** | **AdiC** | **LeuT** |
|---|---|---|---|---|---|---|---|---|---|---|
| | Residues | | | | | | | | | |
| chain A | 2-128 | 2-92 | 22-124 | 17-460 | 3-386 | 2-226 | 29-280 | 12-501 | 5-441 | 13-507 |
| chain B | 5-128 | 2-92 | 22-124 | 18-458 | 3-386 | 2-226 | 29-280 | 12-501 | 5-441 | 13-507 |
| chain C | --- | --- | 22-124 | --- | --- | 2-226 | 29-276 | --- | --- | --- |
| chain D | --- | --- | 22-124 | --- | --- | 2-226 | 29-276 | --- | --- | --- |
| chain E | --- | --- | --- | --- | --- | --- | 30-278 | --- | --- | --- |
| | Number of molecules or ions | | | | | | | | | |
| POPC | 924 | 634 | 609 | 559 | 559 | 551 | 514 | 544 | 567 | 544 |
| $Na^+$ | 90 | --- | --- | 117 | --- | --- | --- | 139 | --- | 135 |
| $K^+$ | --- | 102 | 123 | --- | 135 | 110 | 118 | --- | 120 | --- |
| $Cl^-$ | 98 | 110 | 143 | 134 | 121 | 130 | 124 | 141 | 140 | 145 |
| $H_2O$ | 34163 | 38174 | 45192 | 43231 | 44704 | 39779 | 42997 | 51059 | 43812 | 49662 |
| **System size (nm x nm x nm)** | 15 x 15 x 8.8 | 15 x 15 x 9.7 | 15 x 15 x 9.5 | 15 x 15 x 10.4 | 15 x 15 x 10.6 | 15 x 15 x 9 | 15 x 15 x 10 | 15 x 15 x 11.5 | 15 x 15 x 10 | 15 x 15 x 10 |

**Supplementary Table 5.** System details for oligomeric coarse-grained molecular dynamics simulations – Run 1.

| Run 1 | Fluc-Bpe | Semi SWEET | KcsA | CLC-ec1 | NapA | Aqp0 | FocA | GadC | AdiC | LeuT |
|---|---|---|---|---|---|---|---|---|---|---|
| | Residues | | | | | | | | | |
| chain A | 2-128 | 2-92 | 22-124 | 17-460 | 3-386 | 2-226 | 29-280 | 12-501 | 5-441 | 13-507 |
| chain B | 2-128 | 2-92 | 22-124 | 18-458 | 3-386 | 2-226 | 29-280 | 12-501 | 6-441 | 13-507 |
| chain C | --- | --- | 22-124 | --- | --- | 2-226 | 29-276 | --- | --- | --- |
| chain D | --- | --- | 22-124 | --- | --- | 2-226 | 29-276 | --- | --- | --- |
| chain E | --- | --- | --- | --- | --- | --- | 30-278 | --- | --- | --- |
| Protonated Residues* | GLU88 | --- | GLU71 GLU118 GLU120 | A:GLU113 B:GLU113 B:ASP417 | --- | --- | HIS209 | --- | --- | GLU112 GLU287 GLU419 |
| | Number of molecules or ions | | | | | | | | | |
| POPC | 2002 | 2012 | 1994 | 1092 | 1925 | 1924 | 1877 | 1907 | 1928 | 1901 |
| W | 23495 | 23399 | 23731 | 15218 | 25164 | 25125 | 24905 | 23279 | 23724 | 24929 |
| WF | 2610 | 2599 | 2636 | 1690 | 2796 | 2791 | 2767 | 2586 | 2636 | 2769 |
| $Na^+$ | 283 | 282 | 280 | 178 | 315 | 297 | 305 | 284 | 280 | 300 |
| $Cl^-$ | 291 | 290 | 300 | 195 | 301 | 317 | 305 | 286 | 300 | 310 |
| System size (nm x nm x nm) | 25 x 25 x 10 | 25 x 25 x 10 | 25 x 25 x 10 | 25 x 15 x10 | 25 x 25 x 10 | 25 x 25 x 10 | 25 x 25 x 10 | 25 x 25 x 10 | 25 x 25 x 10 | 25 x 25 x 10 |

*All chains unless otherwise noted.

**Supplementary Table 6.** System details for oligomeric coarse-grained molecular dynamics simulations – Run 2.

| Run 2 | Fluc-Bpe | Semi SWEET | KcsA | CLC-ec1 | NapA | Aqp0 | FocA | GadC | AdiC | LeuT |
|---|---|---|---|---|---|---|---|---|---|---|
| | Residues | | | | | | | | | |
| chain A | 2-128 | 2-92 | 22-124 | 17-460 | 3-386 | 2-226 | 29-280 | 12-501 | 5-441 | 13-507 |
| chain B | 2-128 | 2-92 | 22-124 | 18-458 | 3-386 | 2-226 | 29-280 | 12-501 | 6-441 | 13-507 |
| chain C | --- | --- | 22-124 | --- | --- | 2-226 | 29-276 | --- | --- | --- |
| chain D | --- | --- | 22-124 | --- | --- | 2-226 | 29-276 | --- | --- | --- |
| chain E | --- | --- | --- | --- | --- | --- | 30-278 | --- | --- | --- |
| Protonated Residues* | GLU88 | --- | GLU71 GLU118 GLU120 | A:GLU113 B:GLU113 B:ASP417 | --- | --- | HIS209 | --- | --- | GLU112 GLU287 GLU419 |
| | Number of molecules or ions | | | | | | | | | |
| POPC | 2001 | 2014 | 1993 | 1082 | 1921 | 1921 | 1869 | 1906 | 1921 | 1897 |
| W | 24363 | 25176 | 25054 | 15205 | 23553 | 23474 | 24089 | 23242 | 24992 | 24940 |
| WF | 2707 | 2797 | 2783 | 1689 | 2617 | 2608 | 2676 | 2582 | 2776 | 2771 |
| $Na^+$ | 294 | 304 | 296 | 178 | 295 | 277 | 295 | 283 | 296 | 300 |
| $Cl^-$ | 302 | 312 | 316 | 195 | 281 | 297 | 295 | 285 | 316 | 310 |
| System size (nm x nm x nm) | 25 x 25 x 10 | 25 x 25 x 10 | 25 x 25 x 10 | 25 x 15 x 10 | 25 x 25 x 10 | 25 x 25 x 10 | 25 x 25 x 10 | 25 x 25 x 10 | 25 x 25 x 10 | 25 x 25 x 10 |

*All chains unless otherwise noted.

**Supplementary Table 7.** System details for oligomeric coarse-grained molecular dynamics simulations – Run 3.

| Run 3 | Fluc-Bpe | Semi SWEET | KcsA | CLC-ec1 | NapA | Aqp0 | FocA | GadC | AdiC | LeuT |
|---|---|---|---|---|---|---|---|---|---|---|
| | Residues | | | | | | | | | |
| chain A | 2-128 | 2-92 | 22-124 | 17-460 | 3-386 | 2-226 | 29-280 | 12-501 | 5-441 | 13-507 |
| chain B | 2-128 | 2-92 | 22-124 | 18-458 | 3-386 | 2-226 | 29-280 | 12-501 | 6-441 | 13-507 |
| chain C | --- | --- | 22-124 | --- | --- | 2-226 | 29-276 | --- | --- | --- |
| chain D | --- | --- | 22-124 | --- | --- | 2-226 | 29-276 | --- | --- | --- |
| chain E | --- | --- | --- | --- | --- | --- | 30-278 | --- | --- | --- |
| Protonated Residues* | GLU88 | --- | GLU71 GLU118 GLU120 | A:GLU113 B:GLU113 B:ASP417 | --- | --- | HIS209 | --- | --- | GLU112 GLU287 GLU419 |
| | Number of molecules or ions | | | | | | | | | |
| POPC | 2000 | 1014 | 1991 | 1085 | 1919 | 1919 | 1872 | 1904 | 1924 | 1899 |
| W | 23493 | 23808 | 25027 | 14467 | 23521 | 24291 | 23361 | 24867 | 24189 | 24915 |
| WF | 2610 | 2645 | 2780 | 1607 | 2613 | 2699 | 2595 | 2763 | 2687 | 2768 |
| $Na^+$ | 283 | 287 | 296 | 168 | 295 | 287 | 286 | 303 | 286 | 300 |
| $Cl^-$ | 291 | 295 | 316 | 185 | 281 | 307 | 286 | 305 | 306 | 310 |
| System size (nm x nm x nm) | 25 x 25 x 10 | 25 x 25 x 10 | 25 x 25 x 10 | 25 x 15 x 10 | 25 x 25 x 10 | 25 x 25 x 10 | 25 x 25 X 10 | 25 x 25 x 10 | 25 x 25 x 10 | 25 x 25 x 10 |

*All chains unless otherwise noted.

**Supplementary Table 8.** System details for monomeric coarse-grained molecular dynamics simulations – Run 1.

| **Run 1** | **Fluc-Bpe** | **Semi SWEET** | **KcsA** | **CLC-ec1** | **NapA** | **Aqp0** | **FocA** | **GadC** | **AdiC** | **LeuT** |
|---|---|---|---|---|---|---|---|---|---|---|
| | Residues | | | | | | | | | |
| chain A | 2-128 | 2-92 | 22-124 | 17-460 | 3-386 | 2-226 | 29-280 | 12-501 | 5-441 | 13-507 |
| Protonated Residues | GLU88 | --- | GLU71 GLU118 GLU120 | GLU113 | ASP157 | --- | HIS209 | --- | --- | GLU112 GLU287 GLU419 |
| | Number of molecules or ions | | | | | | | | | |
| POPC | 696 | 703 | 705 | 653 | 659 | 685 | 685 | 1278 | 1285 | 1278 |
| W | 8706 | 8811 | 8814 | 9291 | 9290 | 8711 | 8745 | 15237 | 15174 | 16149 |
| WF | 967 | 979 | 979 | 1032 | 1032 | 967 | 971 | 1693 | 1686 | 1794 |
| $Na^+$ | 105 | 106 | 105 | 109 | 117 | 104 | 106 | 186 | 181 | 196 |
| $Cl^-$ | 109 | 110 | 110 | 118 | 110 | 109 | 107 | 187 | 191 | 200 |
| System size (nm x nm x nm) | 15 x 15 x 10 | 15 x 15 x 10 | 15 x 15 x 10 | 15 x 15 x 10 | 15 x 15 x 10 | 15 x 15 x 10 | 15 x 15 x 10 | 20 x 20 x 10 | 20 x 20 x 10 | 20 x 20 X 10 |

**Supplementary Table 9.** System details for monomeric coarse-grained molecular dynamics simulations – Run 2.

| Run 2 | Fluc-Bpe | Semi SWEET | KcsA | CLC-ec1 | NapA | Aqp0 | FocA | GadC | AdiC | LeuT |
|---|---|---|---|---|---|---|---|---|---|---|
| | Residues | | | | | | | | | |
| chain A | 2-128 | 2-92 | 22-124 | 17-460 | 3-386 | 2-226 | 29-280 | 12-501 | 5-441 | 13-507 |
| Protonated Residues | GLU88 | --- | GLU71 GLU118 GLU120 | GLU113 | ASP157 | --- | HIS209 | --- | --- | GLU112 GLU287 GLU419 |
| | Number of molecules or ions | | | | | | | | | |
| POPC | 692 | 703 | 703 | 646 | 658 | 1313 | 684 | 1279 | 1285 | 1273 |
| W | 8713 | 9314 | 8631 | 9262 | 8727 | 16266 | 8951 | 16109 | 15093 | 16137 |
| WF | 968 | 1034 | 959 | 1029 | 969 | 1807 | 994 | 1789 | 1677 | 1793 |
| $Na^+$ | 105 | 112 | 103 | 109 | 110 | 196 | 109 | 197 | 180 | 195 |
| $Cl^-$ | 109 | 116 | 108 | 118 | 103 | 201 | 110 | 198 | 190 | 200 |
| System size (nm x nm x nm) | 15 x 15 x 10 | 15 x 15 x 10 | 15 x 15 x 10 | 15 x 15 x 10 | 15 x 15 x 10 | 20 x 20 x 10 | 15 x 15 x 10 | 20 x 20 x 10 | 20 x 20 x 10 | 20 x 20 x 10 |

**Supplementary Table 10.** System details for monomeric coarse-grained molecular dynamics simulations – Run 3.

| Run 3 | Fluc-Bpe | Semi SWEET | KcsA | CLC-ec1 | NapA | Aqp0 | FocA | GadC | AdiC | LeuT |
|---|---|---|---|---|---|---|---|---|---|---|
| | Residues | | | | | | | | | |
| chain A | 2-128 | 2-92 | 22-124 | 17-460 | 3-386 | 2-226 | 29-280 | 12-501 | 5-441 | 13-507 |
| Protonated Residues | GLU88 | --- | GLU71 GLU118 GLU120 | GLU113 | ASP157 | --- | HIS209 | --- | --- | GLU112 GLU287 GLU419 |
| | Number of molecules or ions | | | | | | | | | |
| POPC | 695 | 704 | 703 | 650 | 660 | 1312 | 682 | 1278 | 1283 | 1273 |
| W | 9020 | 8793 | 8793 | 9273 | 8729 | 16257 | 8649 | 15274 | 15064 | 16130 |
| WF | 1002 | 977 | 977 | 1030 | 969 | 1806 | 961 | 1697 | 1673 | 1792 |
| $Na^+$ | 108 | 106 | 105 | 109 | 110 | 196 | 105 | 186 | 179 | 195 |
| $Cl^-$ | 112 | 110 | 110 | 118 | 103 | 201 | 106 | 187 | 189 | 200 |
| System size (nm x nm x nm) | 15 x 15 x 10 | 15 x 15 x 10 | 15 x 15 x 10 | 15 x 15 x 10 | 15 x 15 x 10 | 20 x 20 x 10 | 15 x 15 x 10 | 20 x 20 x 10 | 20 x 20 x 10 | 20 x 20 x 10 |
